# Rad51-mediated interhomolog recombination during budding yeast meiosis is promoted by the meiotic recombination checkpoint and the conserved Pif1 helicase

**DOI:** 10.1101/2022.09.05.506590

**Authors:** Andrew Ziesel, Qixuan Weng, Jasvinder S. Ahuja, Abhishek Bhattacharya, Raunak Dutta, Evan Cheng, G. Valentin Börner, Michael Lichten, Nancy M. Hollingsworth

**Author notes:** Corresponding Author: Nancy M. Hollingsworth.

## Abstract

During meiosis, recombination between homologous chromosomes (homologs) generates crossovers that promote proper segregation at the first meiotic division. Recombination is initiated by Spo11-catalyzed double strand breaks (DSBs). 5’ end resection of the DSBs creates 3’ single strand tails that two recombinases, Rad51 and Dmc1, bind to form presynaptic filaments that search for homology, mediate strand invasion and generate displacement loops (D-loops). D-loop processing then forms crossover and non-crossover recombinants. Meiotic recombination occurs in two temporally distinct phases. During Phase 1, Rad51 is inhibited and Dmc1 mediates the interhomolog recombination that promotes homolog synapsis. In Phase 2, Rad51 becomes active and functions with Rad54 to repair residual DSBs, making increasing use of sister chromatids. The transition from Phase 1 to Phase 2 is controlled by the meiotic recombination checkpoint through the meiosis-specific effector kinase Mek1. This work shows that constitutive activation of Rad51 in Phase 1 results in a subset of DSBs being repaired by a Rad51-mediated interhomolog recombination pathway that is distinct from that of Dmc1. Strand invasion intermediates generated by Rad51 require more time to be processed into recombinants, resulting in a meiotic recombination checkpoint delay in prophase I. Without the checkpoint, Rad51-generated intermediates are more likely to be repaired using a sister chromatid, thereby increasing Meiosis I chromosome nondisjunction. This Rad51 interhomolog recombination pathway is specifically promoted by the conserved 5’-3’ helicase *PIF1* and its paralog, *RRM3* and requires Pif1 helicase activity and its interaction with PCNA. This work demonstrates that (1) inhibition of Rad51 during Phase 1 is important to prevent competition with Dmc1 for DSB repair, (2) Rad51-mediated meiotic recombination intermediates are initially processed differently than those made by Dmc1, (3) the meiotic recombination checkpoint provides time during prophase 1 for processing of Rad51-generated recombination intermediates.

**AUTHOR SUMMARY:** To sexually reproduce, cells containing two copies of each chromosome must undergo the specialized cell division of meiosis to sort the chromosomes into gametes containing a single copy of each chromosome. But how do homologous chromosomes know who is who? The answer is by recombination, a process in which double strand breaks on one chromosome are converted to single stranded ends that can search for the complementary sequence on the homolog. In yeast and mammals, this homology search involves binding of single strand ends by two highly conserved recombinases, Rad51 and the meiosis specific Dmc1. Rad51 is used in mitotic cells to repair breaks, primarily using sister chromatids as templates, while Dmc1 functions in meiosis to generate interhomolog crossovers. In budding yeast, Rad51 strand exchange activity is normally inhibited while Dmc1 is active. We show here that when Rad51 and Dmc1 are active at the same time, Rad51 competes with Dmc1 to mediate interhomolog recombination of a subset of double strand breaks. However, because Rad51- generated recombination intermediates take longer to process, there is a need to keep Rad51 inactive while interhomolog recombination is occurring.

## INTRODUCTION

Sexually reproducing diploid organisms create haploid gametes through the specialized cell division of meiosis so that the fusion of two gametes maintains a constant chromosome number in each generation. Meiosis divides the chromosome number in half by having one round of DNA replication followed by two rounds of chromosome segregation. At Meiosis I (MI), homologs segregate to opposite poles, while sister chromatids are disjoined at Meiosis II (MII). Accurately sorting the chromosomes such that each gamete contains exactly one copy requires that homologs be physically connected by a combination of crossovers and sister chromatid cohesion [1]. The process of meiosis is evolutionarily conserved and defects in crossover formation result in increased MI nondisjunction and inviable progeny in organisms ranging from yeast to mammals [2, 3].

The process of crossover formation begins after DNA replication when the highly conserved, meiosis-specific, topoisomerase-like Spo11 protein catalyzes programmed double strand breaks (DSBs) at preferred sites in the genome called hotspots [4–7]. The 5’ ends are then resected, producing 3’ single strand (ss) tails [8–10]. In many organisms, including budding yeast and mammals, these 3’ ends are bound by two recombinases: Rad51, which is also used for mitotic recombination, and the meiosis-specific Dmc1 [11–18]. In yeast and mammals, Rad51 and Dmc1 assemble on the ssDNA in adjacent patches as a right-handed helix to make a presynaptic filament [19–24]. The presence of Rad51, but not its strand exchange activity, is required for normal loading of Dmc1 onto the presynaptic filament and wild-type (WT) levels of meiotic recombination [25–30]. In mice, Dmc1 is localized at the 3’ ends of each filament [22]. The function of recombinases is to mediate a homology search of the genome followed by strand invasion of the donor duplex to form a three-strand structure called a displacement (D) loop [17, 31, 32].

D-loop formation begins when a presynaptic filament interacts with the donor duplex to form a nascent D-loop with an unstable paranemic joint [33–36]. Paranemic joints are subsequently converted to stable plectonemic joints (or mature D-loops) when the recombinase is removed from the invading ssDNA in concert with formation of heteroduplex DNA between the invading strand and the donor strand of opposite polarity, thereby displacing the strand of like-polarity [34, 36, 37]. The bacterial recombinase, RecA, can mediate both the homology search and conversion of nascent to mature D-loops [38]. In contrast, Rad51 can create nascent D-loops, but requires a DNA translocase called Rad54 to make mature D-loops [32, 35, 39, 40].

Rad54 is related to the Snf2 family of helicases and functions as a motor protein that translocates along double-stranded DNA [41, 42]. Rad54 has several functions involved in making D-loops: it directly interacts with Rad51 and stabilizes the Rad51- ssDNA filament, facilitates binding of the presynaptic filament to the donor DNA, and has been proposed to promote a translocation-based homology search along the DNA [39, 43–49]. In addition, Rad54 can remove Rad51 from nascent D-loops leading to a model in which Rad54 acts as a “pump” that couples removal of Rad51 with heteroduplex formation to generate a mature D-loop [37, 50, 51].

Once a mature interhomolog D-loop is made, the 3’ end is extended by DNA Polymerase δ (Pol δ) [52–55]. Some of the extended strands are disassembled from D-loops by the STR (Sgs1, Top3, Rmi1) complex and annealed to the 3’ tails on the other side of the break, leading to noncrossovers through a process called synthesis-dependent strand annealing [56–61]. Alternatively, extension of the invading strand can result in the displaced single strand annealing to the other side of the DSB to create a double Holliday junction [62–64]. During meiosis, most D-loops that form double Holliday junctions are processed by a functionally diverse set of proteins called the “ZMMs” that protect them from disassembly and are resolved almost exclusively as crossovers [56-58, 65, 66]

In addition to promoting interhomolog crossovers, recombination intermediates generated by the ZMM pathway are necessary for chromosome synapsis [66–68]. At the beginning of prophase I, sister chromatids condense along protein cores made of meiosis-specific proteins to create structures called axial elements [69]. In budding yeast, these proteins are Hop1, Red1 and Rec8-containing cohesin [70–72]. ZMM-mediated recombination between homologs creates synaptic initiation sites from which the meiosis-specific transverse filament protein, Zip1, and the central element proteins, Ecm11 and Gmc2, polymerize to “zip” homologous axial elements together to form synaptonemal complexes (SC) [65–67, 73].

A key regulator of meiotic interhomolog recombination is the meiosis-specific kinase Mek1 (also known as Mre4) [74–76]. *MEK1* is required for interhomolog bias, the ZMM crossover pathway and the meiotic recombination checkpoint [77–80]. In mitotic cells, Rad51 mediates DSB repair by homologous recombination with a preference for intersister chromatid recombination [81, 82]. During meiosis, strand invasion is biased towards homologs to create the interhomolog crossovers needed for chromosome disjunction [83]. *MEK1* promotes interhomolog bias in part by inhibiting Rad51-Rad54 complex formation on the presynaptic filament via two mechanisms. First, Mek1 phosphorylation stabilizes the meiosis specific Hed1 protein that binds to Rad51 and excludes Rad54 [84–86]. Second, Mek1 phosphorylation of Rad54 threonine 132 lowers Rad54’s affinity for Rad51 [87]. Mutation of the specific targets of Mek1 that prevent Rad51-Rad54 interaction (*hed1Δ* and/or *RAD54-T132A*) results in efficient DSB repair that is biased towards sister chromatids in a *dmc1Δ* diploid (where Rad51 is the only recombinase), with sufficient interhomolog recombination to produce some viable spores [84, 86, 88–90]. Therefore, allowing Rad51 and Rad54 to interact constitutively during meiosis using the appropriate mutations in *HED1* and *RAD54*, when Mek1 is actively promoting interhomolog bias and Dmc1 is present, results in Rad51-mediated strand invasion of both sister and non-sister chromatids. We refer to this condition as “activated” Rad51.

Mek1 is also the effector kinase for the meiotic recombination checkpoint (MRC) that couples prophase I exit with DSB repair [91]. Mek1 phosphorylation controls the timing of prophase I exit by inhibiting the meiosis-specific transcription factor Ndt80 [77, 91, 92]. Since synapsis only occurs when ZMM crossover-designated recombination intermediates have been generated between homologs, complete chromosome synapsis serves to indicate that additional recombination is no longer necessary [66, 93]. Synapsis reduces Mek1 activity both by decreasing DSB formation and by removing the bulk of Mek1 from chromosomes [94–97]. As a result, Ndt80 is activated and genes required for crossover formation, SC disassembly and meiotic progression are expressed [91, 94, 98–102]. Complete inactivation of Mek1 allows Rad51-Rad54 to repair any remaining DSBs prior to entry into the meiotic divisions [18, 94, 103]. Rad51 activity during meiotic prophase I can therefore be divided into two phases that are delineated by the Mek1-controlled activation state of Ndt80. In Phase 1, Rad51 is inactive and Dmc1 mediates recombination primarily between homologs; in Phase 2, Rad51-Rad54 mediates repair of residual DSBs through strand invasion of sister chromatids. A similar transition from interhomolog to intersister recombination late in prophase I has been observed during nematode meiosis and DSB repair in late meiosis I prophase in mammals also appears to switch to a Rad51-mediated pathway [104–106].

The fact that two independent mechanisms inhibit Rad51 strand invasion in Phase 1 suggests that having Dmc1 and Rad51 active at the same time is disadvantageous to the cell. However, when Rad51 is constitutively activated in the presence of *DMC1* by *hed1Δ* or *hed1Δ RAD54-T132A* (for simplicity, henceforth *hedΔR*), the phenotypes are modest: a two-fold increase in intersister recombination and a delay in meiotic progression with little to no effect on spore viability [88, 89, 107]. Rad51 inhibition is therefore only partially responsible for the interhomolog bias, indicating there are other Mek1 targets yet to be discovered. In contrast, a *hed1Δ dmc1Δ* mutant exhibits an approximately 30-fold decrease in interhomolog bias [88, 89]. The milder phenotypes observed in *hed1Δ* compared to *hed1Δ dmc1Δ* led to the hypothesis that Dmc1 itself partially inhibits Rad51-mediated strand exchange [88, 89]. An alternative hypothesis is that activated Rad51 competes with Dmc1 for strand invasion of DSBs, but that Dmc1 is a more robust recombinase [107]. However, neither of these hypotheses explains why constitutive activation of Rad51 by *hedΔR* results in a meiotic progression delay [88, 107].

In vegetative cells, break-induced replication (BIR) is a DNA repair pathway that occurs when, because homology on the other side of the DSB is poor or absent, the invading strand of a Rad51-mediated D-loop is extended to the end of the chromosome by Polδ [108, 109]. BIR requires the conserved 5’-3’ helicase Pif1 that interacts with PCNA and stimulates Polδ-mediated extension, creating a migrating D-loop [109–111]. Recent work has shown that *PIF1* also has a role in meiotic recombination in yeast [112]. Pif1 localizes to meiotic DSBs, where it interacts with PCNA to promote extension of the invading strand in D-loops formed by Dmc1 [112]. In wild-type (WT) cells, however, the contribution of *PIF1* to strand extension is relatively minor, as its activity is inhibited by the presence of the Mer3-MutLβ complex [112].

We have investigated the basis for the *hedΔR* meiotic progression delay and discovered that allowing Rad51 to interact with Rad54 in Phase 1 results in Rad51 mediated interhomolog strand invasion at a subset of DSBs that is less robust than invasion mediated by Dmc1. The resulting interhomolog recombination intermediates take longer to repair, thereby keeping the MRC active for longer. In the absence of the MRC, constitutive activation of Rad51 results in increased intersister recombination, decreased crossovers, increased MI non-disjunction and reduced spore viability. Furthermore, our work suggests that processing of these intermediates is promoted by *PIF1* and its paralog, *RRM3*. We propose that Rad51 is normally inhibited in Phase 1 to prevent it from competing with Dmc1 for strand invasion and that multiple mechanisms have evolved to promote processing of interhomolog Rad51-mediated D-loops should they occur.

## RESULTS

### Constitutive activation of Rad51 during meiosis delays DSB disappearance and prophase I exit

When Rad51 is active throughout prophase I, entry into the meiotic divisions is delayed [88, 107]. This result was reproduced using the *hedΔR* combination that removes both Mek1-dependent mechanisms that inhibit Rad54 from interacting with Rad51 (Fig 1A). There was significant variability in the kinetics of meiotic progression between different biological replicates of both WT and *hedΔR*, but the amount of sporulation was similar in both strains (Fig 1A-C). To compare the timing of meiotic progression between different strains, a T_50_ value was calculated by determining the time it took for each culture to reach half of its maximum %MI + MII value determined by fluorescent staining of nuclei (MI = binucleate, MII = tetranucleate). The average T_50_ value for *hedΔR* was 6.4 hours, significantly longer than the WT average of 5.6 hours (Fig 1C).

**Fig 1.**
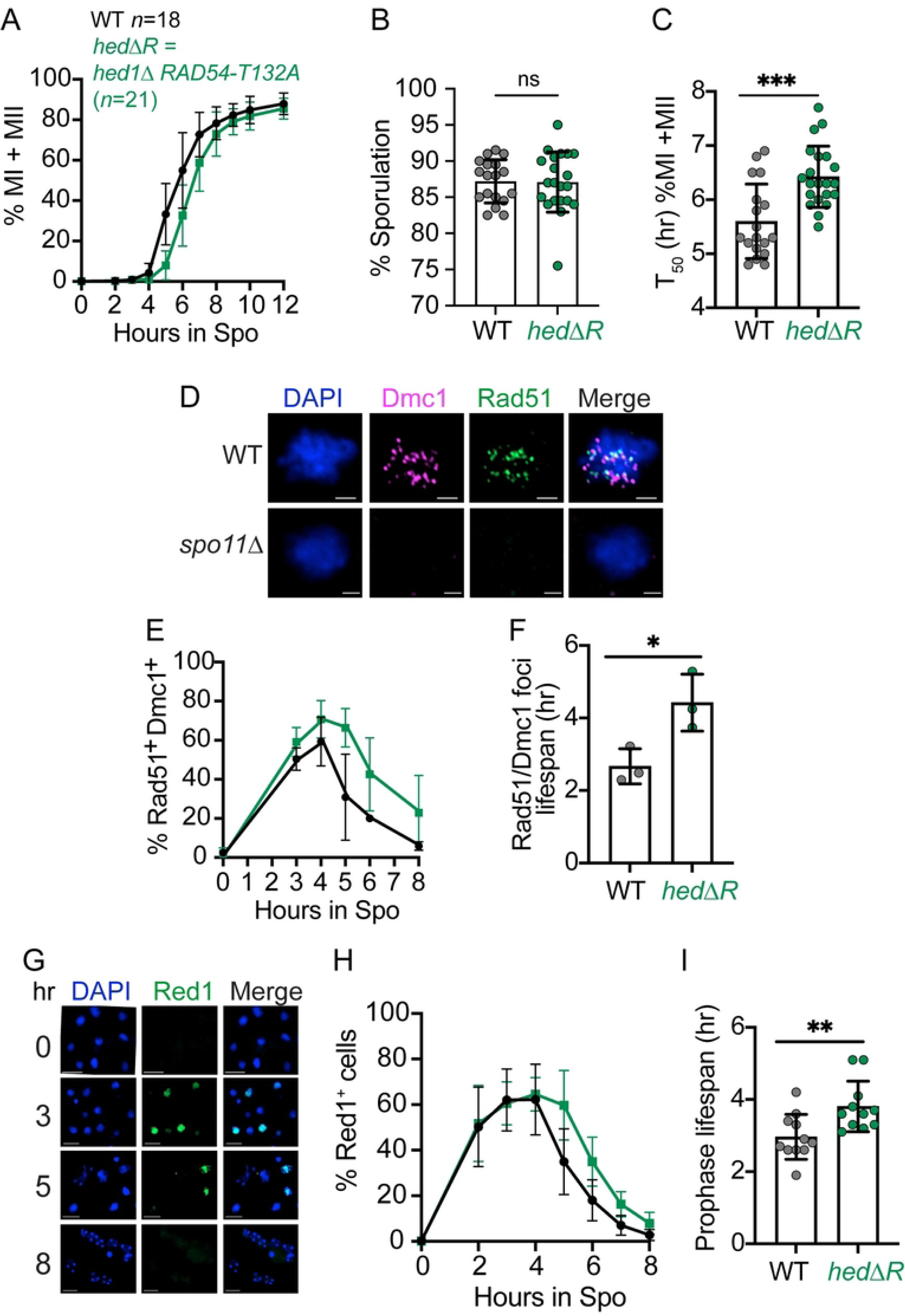
Rad51 activation during Phase 1 causes a delay in meiotic progression, DSB disappearance and prophase I exit. (A) Meiotic progression. WT (NH716 or NH2598 RCEN) (*n*=18) and *hedΔR* (NH2528 or NH2616 RCEN) (*n*=21) diploids were transferred to Spo medium. At the indicated timepoints cells were fixed, stained with 4′,6-diamidino-2-phenylindole (DAPI) and examined by fluorescent microscopy for the presence of binucleate (MI) or tetranucleate (MII) cells. The average values from different timecourses were plotted for each time point. At least 200 cells were counted for each timepoint. (B) Sporulation. Cells were examined by light microscopy for the presence of asci containing one to four spores. For each biological replicate, 200 cells were counted. (C) Quantification of meiotic progression timing. T_50_ represents the amount of time (hr) it took for a sporulation culture from Panel A to reach one-half its maximum %MI+MII value. (D) Cytological assay for the presence of DSBs. Representative images of chromosome spreads generated from WT (NH716) and *spo11Δ* (NH1055) cells taken after four hours in Spo medium. DNA was visualized with DAPI and antibodies against Rad51 and Dmc1 were used to detect the presence of DSBs. Scale bar is 2 µm. (E) Quantification of Dmc1/Rad51 foci. Chromosome spreads of WT (*n*=3), and *hedΔR* (*n*=3) diploids from a subset of the replicates shown in Panel A were analyzed for the presence of Rad51 and Dmc1 foci at various timepoints. Nuclei containing a minimum of three Rad51 and three Dmc1 foci were scored as positive. At least 120 nuclei were examined for each timepoint. (F) Rad51/Dmc1 foci lifespan. Each time course from Panel D was graphed and the lifespan of Rad51/Dmc1 positive nuclei (hr) was calculated by dividing the area under each curve by the maximum number of cells that completed either MI or MII for the corresponding culture [113]. (G) Examples of Red1 whole cell immunofluorescence in a WT meiotic time course. Scale bar is 2 µm. (H) Prophase I progression. Whole fixed cells from a subset of the replicates shown in Panel A (WT and *hedΔR*, *n* = 11) were assayed for the presence of Red1 by immunofluorescence and the average values were plotted. At least 200 cells were counted for each timepoint. (I) Quantification of prophase I length. Prophase I lifespan for each sporulation culture included in H was calculated as described for Panel F. For all bar graphs, each dot represents a biological replicate, the height of the bar indicates the average, and the error bars indicate standard deviation. For all line graphs, error bars indicate standard deviation. Data were analyzed for statistical significance using an unpaired, two-tailed Student T-test. (* = p<.05; ** = p<.01; *** = p<.001)

One explanation for the *hedΔR* meiotic progression delay is that constitutive activation of Rad51 delays DSB repair, thereby keeping the meiotic recombination checkpoint active for longer. DSBs occurring throughout the genome can be cytologically detected in single cells by staining chromosome spreads with antibodies against Rad51 and Dmc1 [27] (Fig 1D). As controls for our antibodies, Rad51 and Dmc1 foci were only observed in meiotic cells, and were dependent on *RAD51* and *DMC1*, respectively (Fig S1A). In addition, few Dmc1 foci were observed in *rad51Δ*, as expected given that *RAD51* is needed to promote assembly of Dmc1 onto the presynaptic filament (Fig S1A) [25, 27]. Finally, no foci were observed in a *spo11Δ* diploid, confirming that the Dmc1/Rad51 foci were dependent on Spo11-generated DSBs (Fig 1D).

In *hedΔR*, cells with both Rad51 and Dmc1 foci accumulated to higher levels and persisted at later timepoints compared to WT (Fig 1E). The lifespan of the period when nuclei have a substantial number of DSBs (≥ 3 Rad51 or Dmc1 foci per nucleus) was determined by graphing the percent of Dmc1 and Rad51 positive spreads at each time point for each replicate, then dividing the area under the curve in each graph by the maximum %MI+MII cells (Fig 1F) [113]. The average Rad51/Dmc1 foci lifespan in *hedΔR* was significantly longer than WT (4.4 vs 2.7 hours) (Note that this analysis underestimates this lifespan in *hedΔR* since the frequency of Rad51^+^ Dmc1^+^ cells did not reach zero by eight hours).

Unrepaired DSBs cause extended activation of the MRC and thus delay exit from prophase I [77, 114, 115]. Measuring meiotic progression by counting the number of nuclei does not provide information as to which part of meiosis is delayed. To determine whether prophase I length is specifically increased by *hedΔR*, whole cell immunofluorescence monitoring the presence of the axial element protein Red1 was used (Fig 1G). Red1 is degraded upon Ndt80 activation due to the appearance of the polo-like kinase, Cdc5, making it an excellent marker for prophase I [94, 98, 101, 103]. The cytological specificity of the Red1 antibody was confirmed by the lack of signal in cells from an *ndt80Δ red1Δ* control (Fig S1B).

In WT meiotic timecourses, the frequency of Red1^+^ cells peaked at three hours and gradually decreased until approaching zero by ∼8 hours (Fig 1H). In *hedΔR*, Red1^+^ cells appeared with WT kinetics but disappeared more slowly than WT. A quantitative value for prophase I lifespan was determined for each replicate by calculating the area under the curve for % Red1^+^ cells and normalizing to the maximum percentage of MI+MII cells. The average prophase I lifespan for *hedΔR* was 3.8 hours, significantly longer than WT (3.0 hours) (Fig 1I). The meiotic progression delay observed for *hedΔR* can be accounted for by the increase in prophase I length, as the difference between *hedΔR* and WT for T_50_ and prophase I lifespan was the same (∼0.8 hours).

### Rad51 strand exchange activity is responsible for the *hed****Δ****R* delay in prophase I exit

Rad51:Dmc1 stoichiometry in the presynaptic filament is important for normal interhomolog recombination [116]. Hed1 and Rad54 can stabilize Rad51 on the filament, raising the possibility that the problems in DNA repair observed in *hedΔR* are due to improperly assembled presynaptic filaments resulting from the absence of *HED1* and/or the presence of Rad54 [49, 117, 118]. An alternative possibility is that presynaptic filament assembly is normal, but Rad51-mediated strand invasion events take longer to process than those created by Dmc1. The latter hypothesis was tested using the *rad51-II3A* mutant, which forms normal presynaptic filaments but is defective in strand exchange [25].

The *rad51-II3A* diploid exhibited a meiotic progression delay and a small, but significant, decrease in spore viability, likely due to the failure of Rad51 to repair residual DSBs in Phase 2 (Fig 2A-C) [25, 94, 103]. Consistent with this idea, the distribution of viable spores in *rad51-II3A* tetrads was consistent with random spore viability and MI nondisjunction was not increased (Fig 3FG). Furthermore, while the T_50_ value for meiotic progression in *rad51-II3A* was significantly greater than in WT, prophase I length was not significantly different (Fig 2CE). One explanation for the *rad51-II3A* meiotic progression delay is that some DSBs persist past MI, thereby triggering the DNA damage checkpoint prior to MII [119].

**Fig 2.**
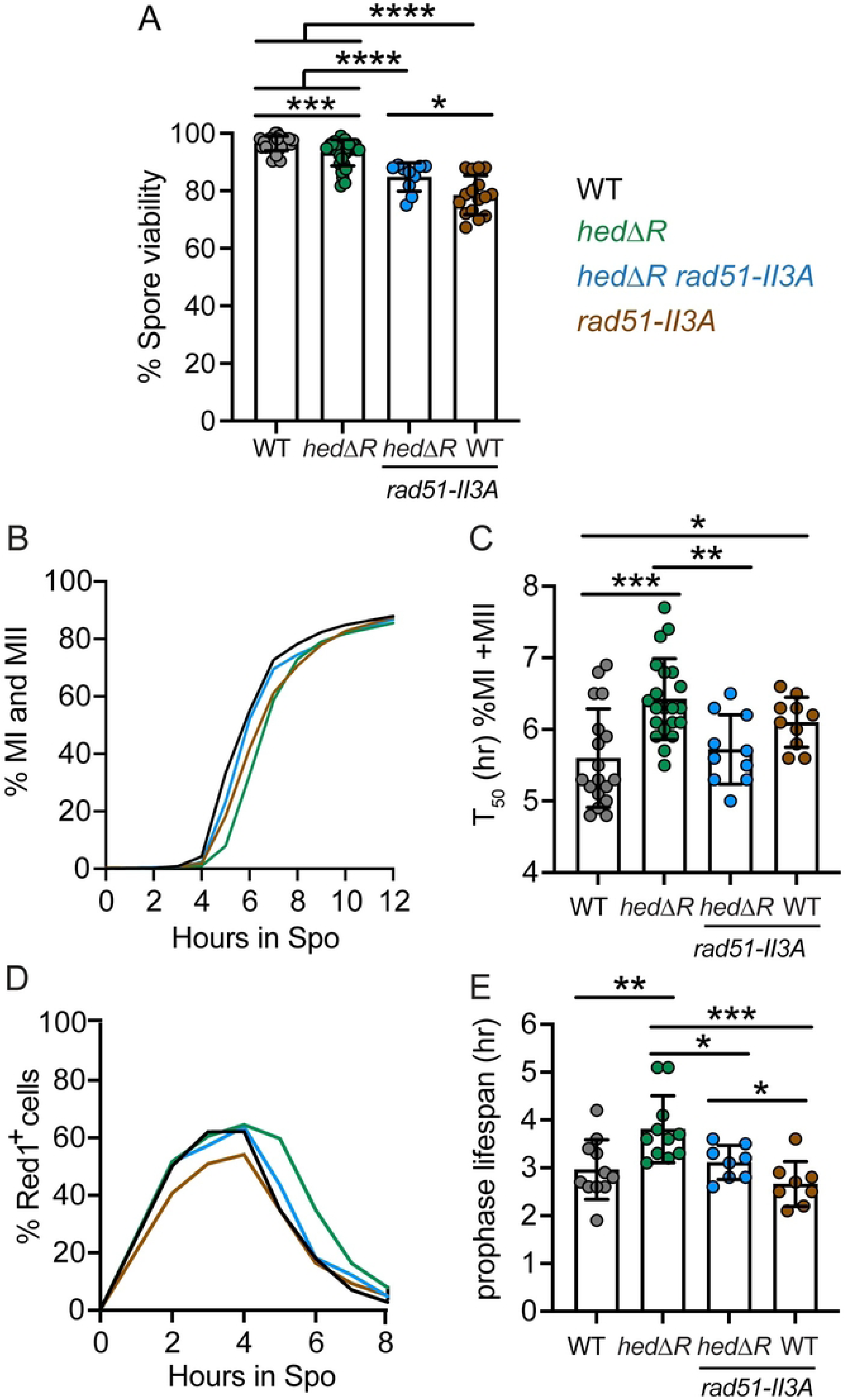
Activation of Rad51-dependent delays prophase I exit. (A) Spore viability. WT (NH716 or NH2598 RCEN), *hedΔR* (NH2528 or NH2616 RCEN), *hedΔR rad51-II3A* (NH2566) and *rad51-II3A* (NH2618 or NH2666 RCEN) tetrads from both liquid and solid sporulation conditions were dissected to determine the percent of viable spores. For each replicate, at least 18 tetrads were dissected. (B) Meiotic progression. Timecourses of the WT, *hedΔR*, *hedΔR rad51-II3A* (*n*=10) and *rad51-II3A* (*n*=10) were analyzed for meiotic progression as described in Figure 1. (C) Comparison of T_50_ values of the individual timecourses shown in Panel B. (D) Prophase I progression. A subset of timecourses in panel A (WT and *hedΔR*, *n*=11, *rad51-II3A* and *hedΔR rad51-II3A*, *n*=8) were analyzed for prophase I progression as described in Figure 1. (E) Prophase I lifespan for each sporulation culture in D was calculated as described in Figure 1. The data for WT and *hedΔR* used for Panels B and D are the same as shown in Figure 1. The statistical significance of differences between strains in Panel A used the Mann-Whitney test, while Panels C and E used an unpaired, two-tailed Student’s T-test. (* = *p*<.05; ** = *p*<.01; *** = *p*<.001, **** = *p*<.0001).

**Fig 3.**
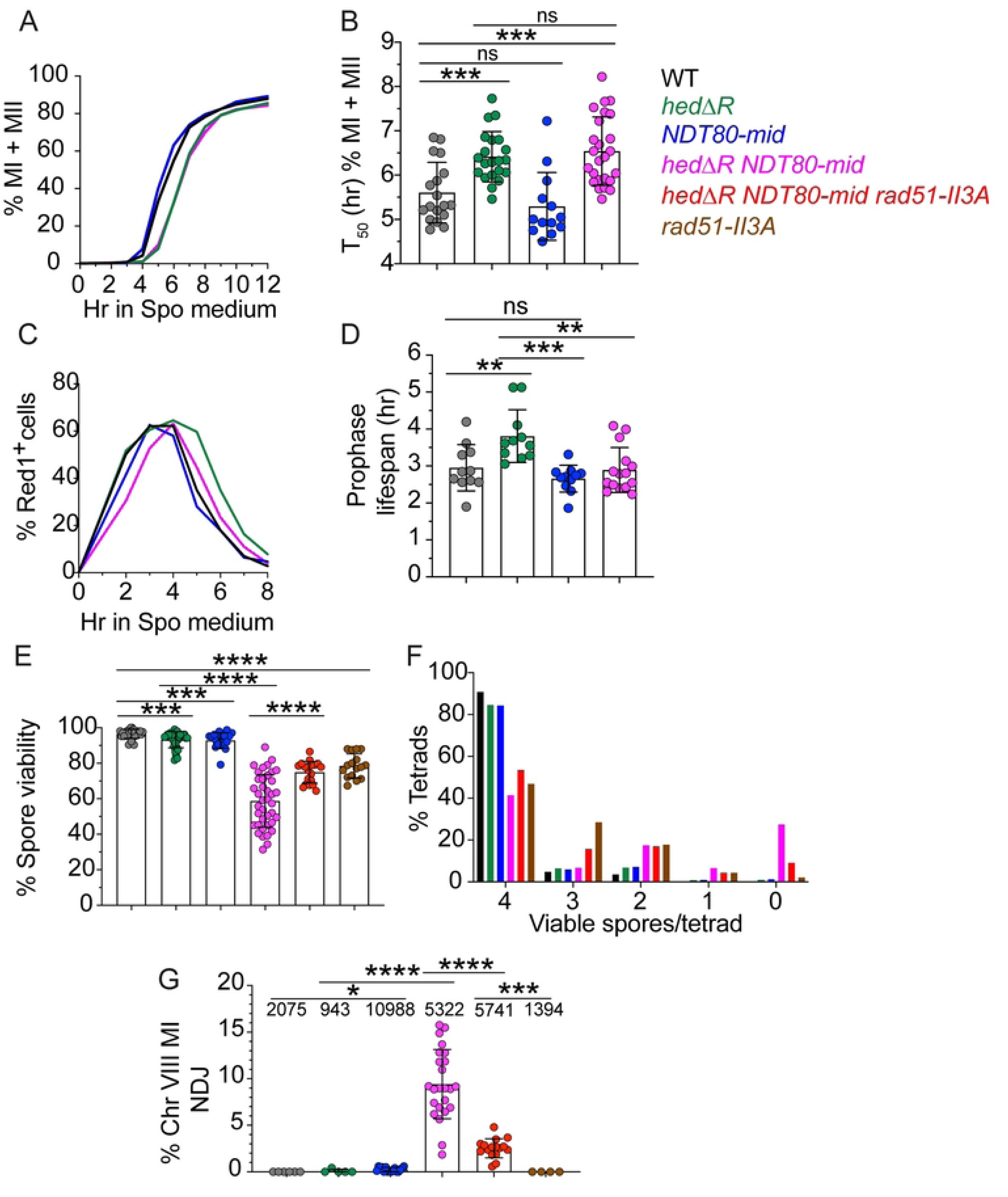
Phenotypic consequences of constitutively activating Rad51 in the absence of the meiotic recombination checkpoint. (A) Meiotic progression. Timecourses of the following diploids were analyzed for meiotic progression as described in Figure 1: WT, *hedΔR*, *NDT80-mid* (NH2495 or NH2634 RCEN) (*n*=13) and *hedΔR NDT80-mid* (NH2505 or NH2610 RCEN) (*n*=25). (B) Comparison of T_50_ values of timecourses shown in Panel A. (C) Prophase I progression. A subset of timecourses in panel A (*NDT80-mid*, *n* =11, *hedΔR NDT80-mid*, *n* =15) were analyzed for prophase I progression as described in Fig 1. WT and *hedΔR* data are from Fig 1. (D) Prophase I lifespan for each sporulation culture in C was calculated as described in Fig 1. (E) Spore viability. Sporulated cells from both liquid and solid sporulation media were dissected to determine the percent of viable spores. At least 20 tetrads were dissected for each replicate. In addition to the diploids listed for Panel A, *hedΔR NDT80-mid rad51-II3A* (NH2705 RCEN) and *rad51-II3A* were included. (F) The distribution of viable spores among tetrads dissected in Panel E. (G) Frequency of chromosome VIII nondisjunction during MI. Numbers above each bar indicate the total number of tetrads assayed. Panels A-E for WT, *hedΔR*, and *rad51-II3A* used the same data shown in Figures 1 and 2. The statistical significance of differences between strains in Panels B and D was determined using an unpaired, two-tailed Student’s T-test, while the Mann-Whitney test was used for Panels E and G. (* = *p*<.05; ** = *p*<.01; *** = *p*<.001, **** = *p*<.0001)

To see if constitutive Rad51-mediated strand invasion delays prophase I exit, *rad51-II3A* was combined with *hedΔR*, which should allow the strand exchange defective Rad51-II3A protein to interact with Rad54. The *hedΔR rad51-II3A* diploid exhibited lower spore viability than *hedΔR*, as expected since the constitutively active Rad51-II3A protein is unable to repair residual DSBs during Phase 2 (Fig 2A). In contrast, preventing Rad51 mediated strand invasion in the *hedΔR rad51-II3A* diploid suppressed the *hedΔR* meiotic progression delay by decreasing the length of prophase I (Fig 2B-E). The average T_50_ value for *hedΔR rad51-II3A* was 5.7 hours, significantly less than the 6.4 hours observed for *hedΔR*. In addition, prophase I lifespan was decreased significantly from 3.8 hours in *hedΔR* to 3.1 hours in *hedΔR rad51-II3A*. These results show that constitutively allowing Rad54 to interact with Rad51 during meiosis results in strand exchange events that trigger a delay during prophase I.

### The prophase I delay caused by constitutive activation of Rad51 depends on the MRC

The delayed exit from prophase I and persistence of DSBs in *hedΔR* suggests that MRC activity is extended in response to activated Rad51. If true, then eliminating this checkpoint should rescue the *hedΔR* prophase I delay. Mek1 controls the MRC by binding to an RPSKR motif in Ndt80 and phosphorylating the Ndt80 DNA binding [91]. An *NDT80* allele lacking the RPSKR motif (called *NDT80-mid*, for <u>M</u>ek1 Interaction <u>D</u>efective) makes Ndt80 insensitive to the checkpoint without affecting its ability to activate transcription [91].

The *NDT80-mid* diploid exhibited similar meiotic progression timing as WT (Fig 3AB). However, abrogating the checkpoint in the presence of activated Rad51 (*hedΔR NDT80-mid*) did not eliminate the meiotic progression delay seen in *hedΔR* (Fig 3A, B). One explanation for the failure of *NDT80-mid* to suppress the *hedΔR* meiotic progression delay is that this mutant combination lengthens meiotic progression for a different reason than *hedΔR* alone: e.g., a delay in the metaphase to anaphase transition due to activation of the spindle assembly checkpoint in response to nonexchange chromosomes [120]. Since the MRC specifically controls the timing of prophase I exit, prophase I lifespan was determined for *hedΔR NDT80-mid*. For reasons that are unclear, *hedΔR NDT80-mid* was delayed in entering prophase I compared to WT, *NDT80-mid* and *hedΔR* (Fig 3C). Despite entering prophase I later, *hedΔR NDT80- mid* cells exited prophase I significantly earlier than *hedΔR*, with an average lifespan of 2.9 hours compared to 3.8 hours for *hedΔR* (Fig 3D). These data indicate that constitutive activation of Rad51 during meiosis results in strand exchange intermediates that take longer to process and trigger the MRC to delay exit from prophase I.

### The MRC is required for proper Meiosis I disjunction and spore viability when Rad51 is constitutively active during prophase I

If the MRC is providing time for cells to process strand invasion intermediates generated by Rad51, there should be phenotypic consequences when the checkpoint is absent. Diploids in which either Rad51 was constitutively activated (*hedΔR*), or the MRC was eliminated (*NDT80-mid*), exhibited very small, but significant, decreases in spore viability (Fig 3E). In contrast, activating Rad51 in the absence of the checkpoint in the *hedΔR NDT80-mid* diploid dramatically reduced spore viability to an average of ∼60% (Fig 3E). This value is significantly less than the value predicted from the product of the individual spore viabilities of *hedΔR* and *NDT80-mid* (86.5%), indicating a synergistic effect. Furthermore, there was increased variability in the spore viability observed between individual colonies in the *hedΔR NDT80-mid* strain compared to *hedΔR* alone (Fig 3E).

The tetrads from *hedΔR NDT80-mid* exhibited a pattern of spore inviability consistent with MI nondisjunction, where four viable spore tetrads are reduced in favor of two and zero viable spore tetrads (Fig 3F) [121]. To monitor MI disjunction directly, genes encoding either a spore autonomously-expressed red fluorescent protein (*RFP*) or cyan fluorescent protein (*CFP*) were introduced at allelic positions adjacent to the centromere of Chromosome VIII (Fig S2A) [122]. Proper segregation of Chromosome VIII produces two red and two blue spores in each tetrad, while MI non-disjunction results in two pink and two black spores (Fig S2A). No MI nondisjunction was observed out of 2075 WT tetrads (Fig 3G). Like WT, chromosome segregation was highly accurate in the *hedΔR* diploid, while a small, but significant, increase in MI nondisjunction was detected for *NDT80-mid* (Fig 3G). MI nondisjunction was dramatically increased in the *hedΔR NDT80-mid* strain, consistent with the observed decrease in spore viability (Fig 3G).

The observed *hedΔR NDT80-mid* disjunction and spore viability defects are due to Rad51 mediated strand invasion because disrupting Rad51’s strand invasion activity in the *hedΔR NDT80-mid rad51-II3A* diploid restored spore viability to the same level as *rad51-II3A* alone, as well as significantly reducing MI nondisjunction (Fig 3 EG). The MRC therefore has a vital role in promoting proper chromosome segregation when Rad51 is constitutively activated.

### The MRC promotes crossover formation in *hed****Δ****R* diploids

One explanation for the large increase in MI non-disjunction in *hedΔR NDT80- mid* is that crossover formation is reduced. Crossovers were measured in two ways that do not require viable spores. The first method used physical analysis of crossover and noncrossover products resulting from DSB repair of the *HIS4LEU2* hotspot [92, 123]. The DSB site of this meiotic hotspot is flanked with XhoI restriction sites that are differentially located on each homolog, with an NgoMIV site very near the DSB site on one of the homologs (Fig 4A) [123]. Double digestion of the genomic DNA with XhoI and NgoMIV produces unique fragments indicative of a subset of crossovers (CO2) and noncrossovers (NCO1) (Fig 4A). Meiotic timecourse analysis revealed that similar to previous results, both NCO1 and CO2 were delayed and reduced ∼2 fold in *hedΔR* (Fig S3CB)[88, 107]. However no further decrease in the amount of either CO2 or NCO1 was observed in the *hedΔR NDT80-mid* diploid when the average values were plotted for two different timecourses (Fig S3CB).

**Fig 4.**
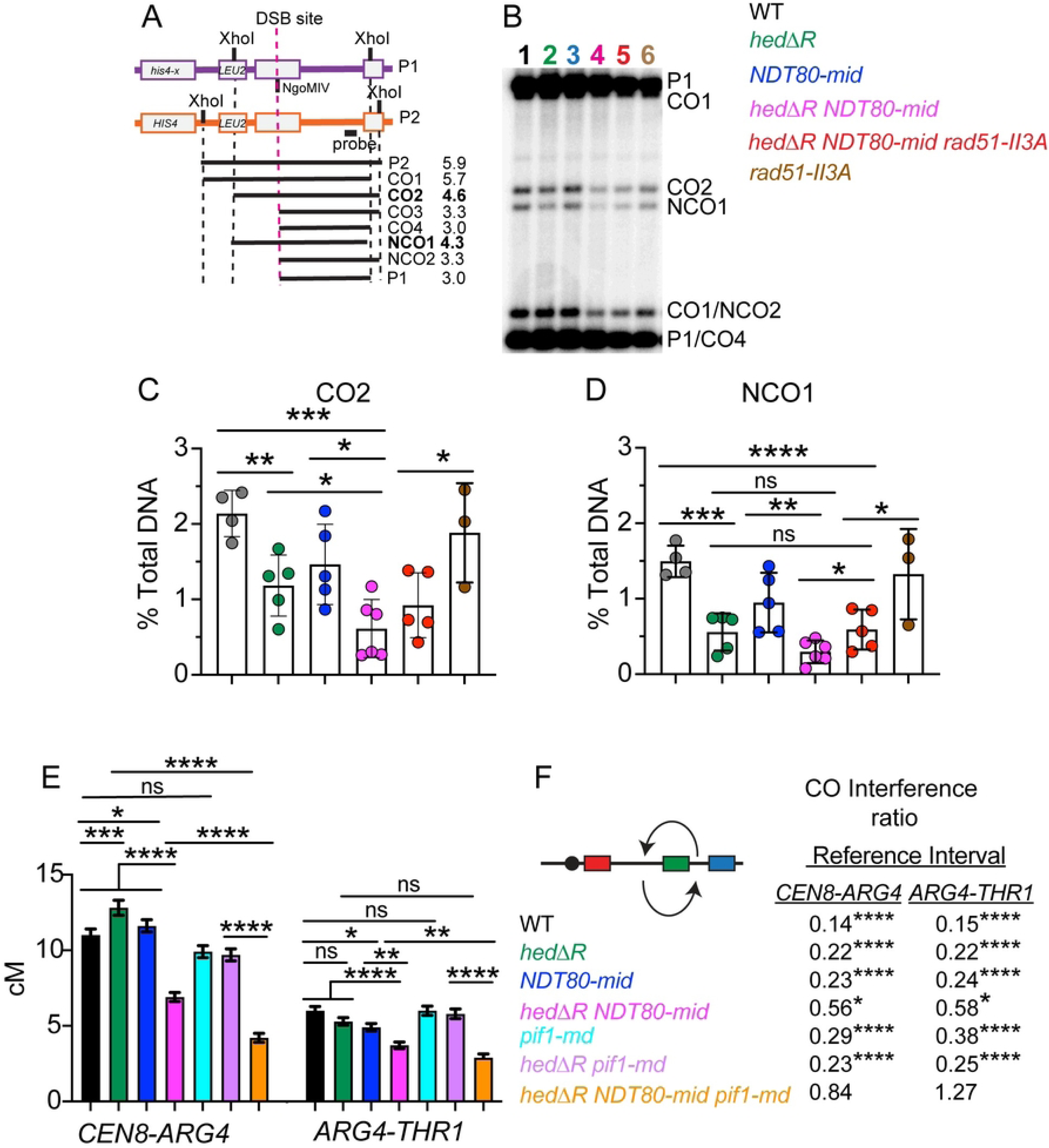
Physical and genetic analysis of recombination in various mutants. (A) Physical analysis of recombination. Schematic of the *HIS4LEU2* hotspot showing the fragments and their respective sizes in kb predicted after digestion with XhoI and NgoMIV (adapted from Wu, Ho (92)). Black dotted lines indicate ends generated by XhoI. The red dotted lines indicate the DSB made by Spo11. NCO1 and CO2 are unique restriction fragments indicated in bold. (B) Cultures generated from independent single colonies of WT (NH716 and NS2598 RCEN), *hed1ΔR* (NH2528 and NH2H2616 RCEN), *NDT80-mid* (NH2634 RCEN), *hedΔR NDT80-mid* (NH2505 and NH2610), *hedΔR NDT80-mid rad51-II3A* (NH2705 RCEN) and *rad51-II3A* (NH2666 RCEN) were incubated for 10 hours in Spo medium. DNA was isolated, digested with XhoI and NgoMIV and crossovers and noncrossovers were analyzed by Southern blot as described in [123]. A representative gel of an individual culture from each strain. Numbers are colored coded with the indicated strains in the legend. (C) Quantification of CO2. (D) Quantification of NCO1. The statistical significance of differences between strains in Panels B and C was determined using an unpaired, two-tailed Student’s T-test. (* = *p*<.05; ** = *p*<.01; *** = *p*<.001, **** = *p*<.0001). (E) Genetic analysis of crossover formation. Single colonies from WT (NH2598 RGC) (*n* = 7; tetrads = 3374), *hedΔR* (NH2616 RGC) (*n* = 7; tetrads = 3812), *NDT80-mid* (NH2634 RGC) (*n* = 7; tetrads = 3592), *hedΔR NDT80-mid* (NH2610 RGC) (*n* = 8; tetrads = 3899), *pif1-md* (NH2695 RGC)(*n* = 7; tetrads scored = 2974), *hedΔR pif1-md* (NH2702 RGC)(*n* = 6; tetrads = 2799) and *hedΔR NDT80-mid pif1-md* (NH2687 RGC)(*n* = 8; tetrads = 2605) were sporulated on solid medium and analyzed by fluorescence microscopy for crossovers in the *CEN8-ARG4* and *ARG4-THR1* intervals (Fig S4). Black bars show the average of plotted dots, bars show standard error. Since MI nondisjunction and nonparental ditype tetrads are indistinguishable in the *CEN8-ARG4* interval (Fig S4), the frequency of nonparental ditypes was corrected using the MI nondisjunction frequency for *hedΔR* and *NDT80-mid*. No correction was necessary for WT because MI nondisjunction was not observed. All *CEN8-ARG4* NPDs were assumed to be due to MI nondisjunction in strains with Chromosome VIII MI nondisjunction frequencies ≥ 9% (Fig 3G) [122]. Map distances (cM) and standard errors were calculated using the Stahl lab online tools (https://elizabethhousworth.com/StahlLabOnlineTools/). (F) Crossover interference ratios between the *CEN8-ARG4* and *ARG4-THR1* intervals were determined using the method described by Malkova, Swanson (137). Arrows above and below the chromosome VIII diagram indicate *ARG4-THR1* and *CEN8-ARG4* as the reference interval, respectively. Test interval/Reference interval map distance ratios significantly less than one are indicative of interference and *p* values are indicated by asterisks (*<0.05; ****<0.0001) (G-test).

Because of the high variability in spore viability exhibited by different biological replicates of *hedΔR NDT80-mid*, more than two replicates are necessary to determine statistical significance. End point analysis was therefore performed by comparing CO2 and NCO1 frequencies in several biological replicates at the 10 hour timepoint, by which time all of the strains had completed either MI or MII (Figure 2B, 3A). CO2 and NCO1 fragments were quantified as a percent of total DNA to allow statistical significance to be determined. With the larger sample size, a statistically significant reduction in the number of crossovers was observed when *hedΔR* was combined with *NDT80-mid*, correlating well with the spore inviability and MI nondisjunction phenotypes (Fig 4BC). Noncrossovers were also reduced two-fold relative to *hedΔR* in the *hedΔR NDT80-mid* diploid, but this reduction was not statistically significant, perhaps due to the limited sample size (Fig 4BD).

The *rad51-II3A* mutant alone exhibited CO2 and NCO1 levels equivalent to that of WT as expected (Fig 4B-D) [25]. Addition of *rad51-II3A* to the *hedΔR NDT80-mid* diploid increased NCO1 levels, but a statistically significant increase in CO2 was not observed compared to *hedΔR NDT80-mid* (Fig 4CD). However, given the suppression of both MI non-disjunction and spore inviability phenotypes in *hedΔR NDT80-mid rad51-II3A* diploids, the failure to see suppression of crossover formation could be due to the limited sample size and the variability in the rate of meiotic progression observed for *hedΔR NDT80-mid*.

Crossover frequencies were also determined genetically using a fluorescent spore assay [122]. One chromosome VIII contained *RFP* adjacent to *CEN8*. On the homolog, a *GFP\** (green fluorescent protein) gene was integrated at *ARG4* and *CFP* was integrated at *THR1* (Fig S2B). Crossovers in different intervals were distinguished by distinctive fluorescent spore patterns (Fig S4). In contrast to the *HIS4LEU2* hotspot analysis, the map distance in the *CEN8-ARG4* interval was higher than WT for both *hedΔR* and *NDT80-mid* (Fig 4E). This increase was specific to the centromere adjacent region, as *hedΔR* and *NDT80-mid* crossovers in the *ARG4-THR1* interval occurred at frequencies that were either equal to or lower than WT (Fig 4E). Interestingly a similar pattern was observed for map distances in different intervals of chromosome III in *hed1Δ* and WT diploids [88]. In that experiment, crossover formation was higher in the *CEN3-MAT* interval in the *hed1Δ* deletion but equal to or lower than WT in intervals further out on the arms. Eliminating the MRC in the Rad51-activated strain (*hedΔR NDT80-mid*) significantly decreased crossovers for both the *CEN8-ARG4* and *ARG4-THR1* intervals compared to *hedΔR* (Fig 4E). These results indicate that a subset of interhomolog recombination intermediates created when Rad51 is activated in Phase 1 require the MRC to be successfully processed into crossovers.

### Constitutive activation of Rad51 results in an accumulation of interhomolog and intersister joint molecules that is dependent upon the MRC

Crossovers are the end point of recombination formed by the resolution of double Holliday junction intermediates (dHJs). Previous work using *hed1Δ* showed a modest decrease in interhomolog (IH) dHJs with little to no effect on intersister (IS) dHJs and a reduction in interhomolog bias from 5.7 IH:IS ratio to 2.1 [88]. In our *hedΔR* diploid, the peak level of IH dHJs was delayed relative to WT, indicating a delay either in strand invasion or second end capture (Fig 5AB). IS-dHJs were increased in *hedΔR* and the IH:IS ratio bias was reduced from ∼8:1 in WT to 2:1 in *hedΔR* (Fig 5AB, S1_Data). In cells lacking the MRC (*NDT80-mid*), the kinetics of IH and IS dHJs was similar to WT and there was a specific reduction in IH dHJs, decreasing the IH:IS ratio to 5:1 (Fig 5AB). The reduction in IH dHJs in *NDT80-mid* could be due to more rapid resolution of the dHJs since transcription of *CDC5* is no longer regulated. This reduction is consistent with the small, but significant, decrease in COs in the *ARG4-THR1* interval and a small, but significant, increase in chromosome VIII MI non-disjunction (Fig 3G, 4E).

**Fig 5.**
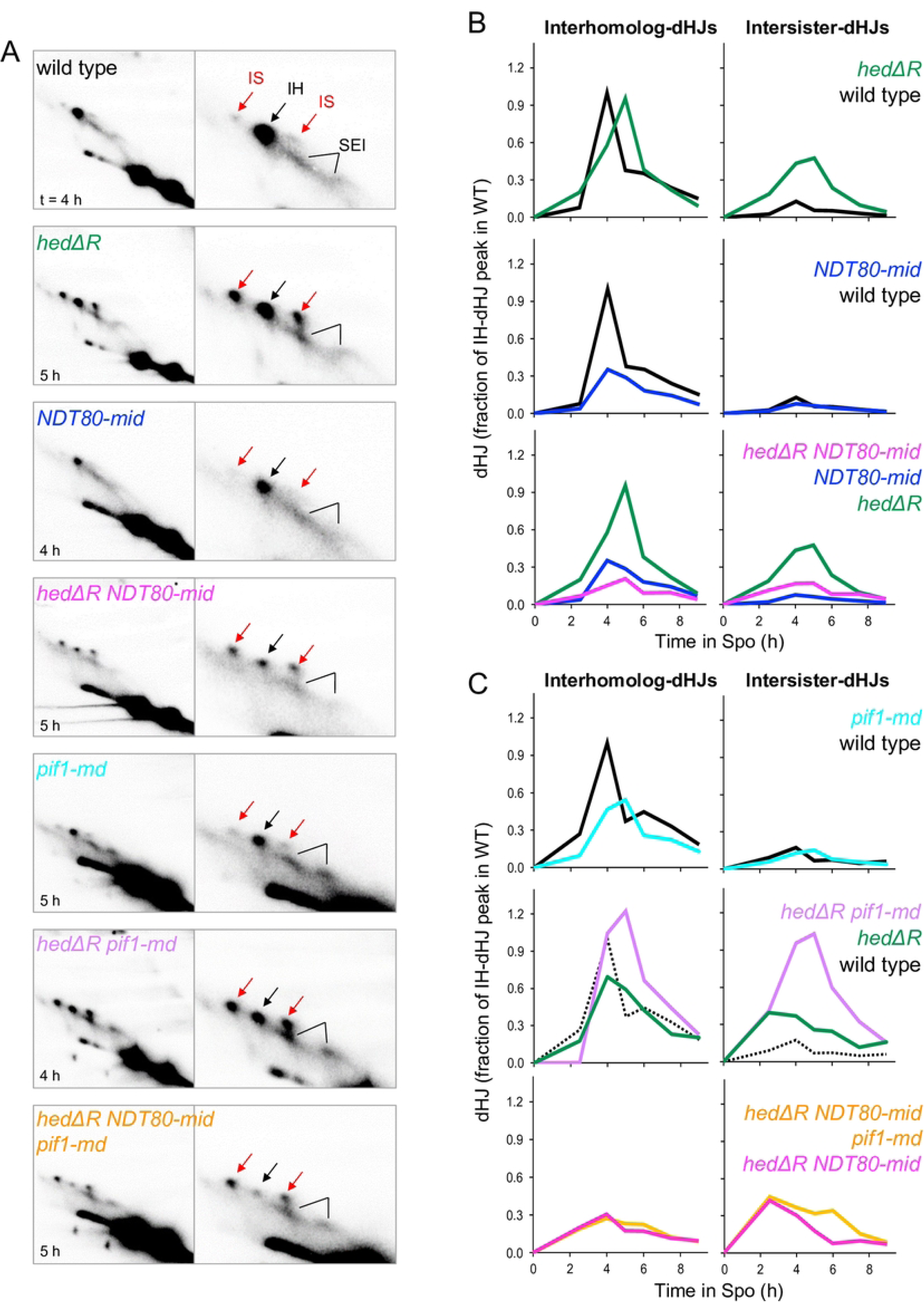
The meiotic recombination checkpoint promotes Rad51-mediated dHJ formation. (A) Excerpts of representative two-dimensional gel Southern blot analyses at the time of maximum joint molecule levels in WT, *hedΔR*, *NDT80-mid*, *hedΔR NDT80-mid*, *pif1-md*, and *hedΔR NDT80-mid pif1-md*. Right panels show enlarged excerpts of left panels. Interhomolog dHJs (black arrow), intersister dHJs (red arrows), interhomolog and intersister SEIs (black lines). (B) Quantitative analysis of effects of *hedΔR* and *NDT80-mid* on IH-dHJs (left) and IS-dHJs (right; time course tc17-1). Joint molecule (JM) levels are expressed as fraction of the maximum IH-dHJ levels in the parallel WT culture (black; here 1.7% of total hybridization signal). (C) Quantitative analysis of effects of *pif1-md* in the WT, *hedΔR* and *hedΔR NDT80-mid* backgrounds on IH-HJs (left) and IS-dHJs (right; tc14). JM levels are expressed as fraction of the maximum IH-dHJ levels in the parallel WT culture (black), here 1.0% of total hybridization signal.

Activating Rad51 in the absence of the MRC (*hedΔR NDT80-mid*) greatly reduced both IH and IS dHJs compared to *hedΔR* and the IH:IS ratio was decreased to 1:1 (Fig 5AB, S1_Data). Although there were lower steady state levels of IS dHJs than in *hedΔR*, this amount was increased relative to *NDT80-mid*, potentially explaining the decrease in crossovers observed when Rad51 is activated in the absence of the MRC.

### Extended time in prophase I promotes chromosome disjunction when Rad51 is constitutively activated

An unexpected finding was the amount of variability in prophase I length between cultures derived from different single colonies of the same diploid, including WT (Fig 3D). Despite this variability, there was no correlation between spore viability and prophase I length in either WT, *hedΔR*, or *hedΔR rad51-II3A* (Fig 6A). In addition, specifically eliminating the checkpoint in WT cells using *NDT80-mid* did not change this lack of correlation. In contrast, prophase I length was positively correlated with spore viability and inversely correlated with chromosome VIII MI nondisjunction in *hedΔR NDT80-mid* cultures (Fig 6A, B). The fact that these correlations were lost in the *hedΔR NDT80-mid rad51-II3A* strain indicates that Rad51 strand invasion activity is required for this phenotype (Fig 6A, B). We conclude that the longer prophase I lengths observed for *hedΔR* are due to the creation of recombination intermediates that take longer to be productively processed into crossovers than those mediated by Dmc1. In the absence of the checkpoint, the length of prophase I is no longer controlled, and so cells without a sufficient number of crossovers prematurely enter MI, resulting in homolog nondisjunction.

**Fig 6.**
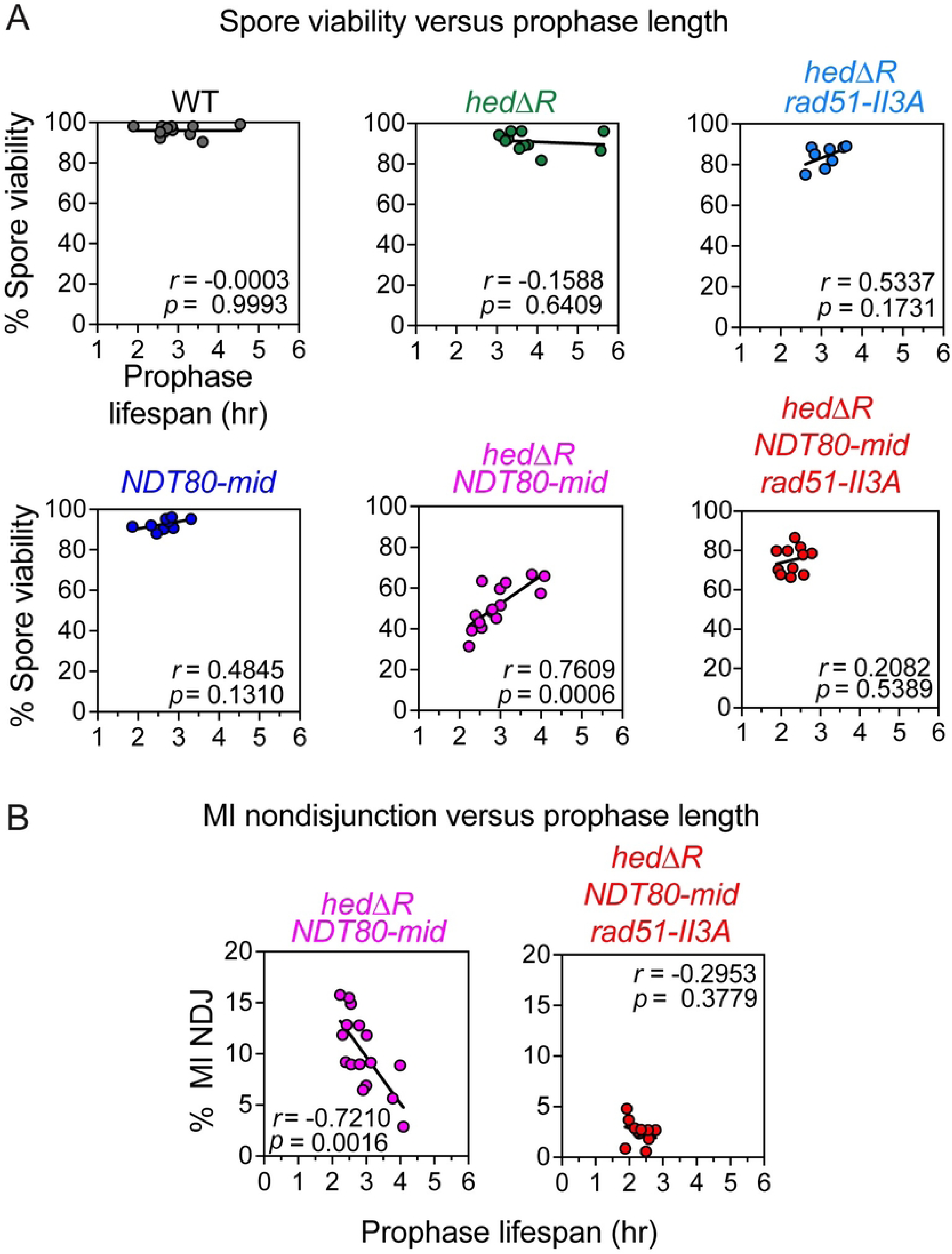
The meiotic recombination checkpoint uncouples prophase I length from spore viability and MI chromosome segregation when Rad51 is constitutively activated. (A) Spore viability was plotted versus prophase I length for WT (*n* = 12), *hedΔR* (*n* = 11), *hedΔR rad51-II3A* (*n* = 8), *NDT80-mid* (*n* = 11), *hedΔR NDT80-mid* (*n* = 16) and *hedΔR NDT80-mid rad51-II3A* (*n* = 11). (B) Chromosome VIII MI nondisjunction was plotted versus prophase I length for *hedΔR NDT80-mid* and *hedΔR NDT80-mid rad51-II3A*, using the same replicates as in Panel A. These data are from the same timecourses shown in Fig 3. The correlation coefficient and statistical significance are indicated by “*r*” and “*p*”, respectively and were determined using GraphPad Prism 9.0.

### Strand invasion is less robust when Rad51 is constitutively active

The persistence of Rad51/Dmc1 foci in strains where Rad51 is constitutively activated, as well as the need for a checkpoint-mediated delay to successfully complete meiotic recombination in *hedΔR* strains, suggests that strand invasion might be less robust than in WT. Previous studies have suggested that meiotic recombination frequently involves multiple rounds of strand invasion and D-loop intermediate disassembly before stable recombination intermediates or products form by strand annealing [124–127]. Less robust strand invasion, by shifting the balance away from D-loop formation and towards disassembly/annealing, might be expected to decrease the number of strand invasions that occur before stable intermediates or products form.

To test this idea, the parental strand contributions to recombinants in a highly polymorphic test interval where recombination is initiated by a tightly-focused DSB at the *URA3-tel-ARG4* hotspot (Fig 7A) were determined using high throughput sequencing to score segregation of the polymorphic markers in 159 tetrads from an *hedΔR msh2Δ* diploid [126]. While crossing-over between *natMX* and *hphMX* inserts flanking the interval was similar in *hedΔR* and WT (26.8 ± 2.6 and 30.4 ± 1.9 cM, respectively; S2_Data, Sheet 13), gene conversion frequencies of markers were reduced in *hedΔR* by about 25% through much of the interval (p <0.0001, Wilcoxon matched-pairs signed rank test; Fig 7B and S2_Data, Sheet 4), consistent with reduced strand invasion. In addition, while the majority of events in WT contained heteroduplex DNA on both sides of the DSB, only a minority of events in *hedΔR* were two-sided (Fig 7C, S5, S2_Data, Sheet 5). Since two-sided events can be produced by events where both DSB ends invade and prime synthesis, the reduced number of such events provides additional indication of reduced strand invasion in *hedΔR*.

**Fig 7.**
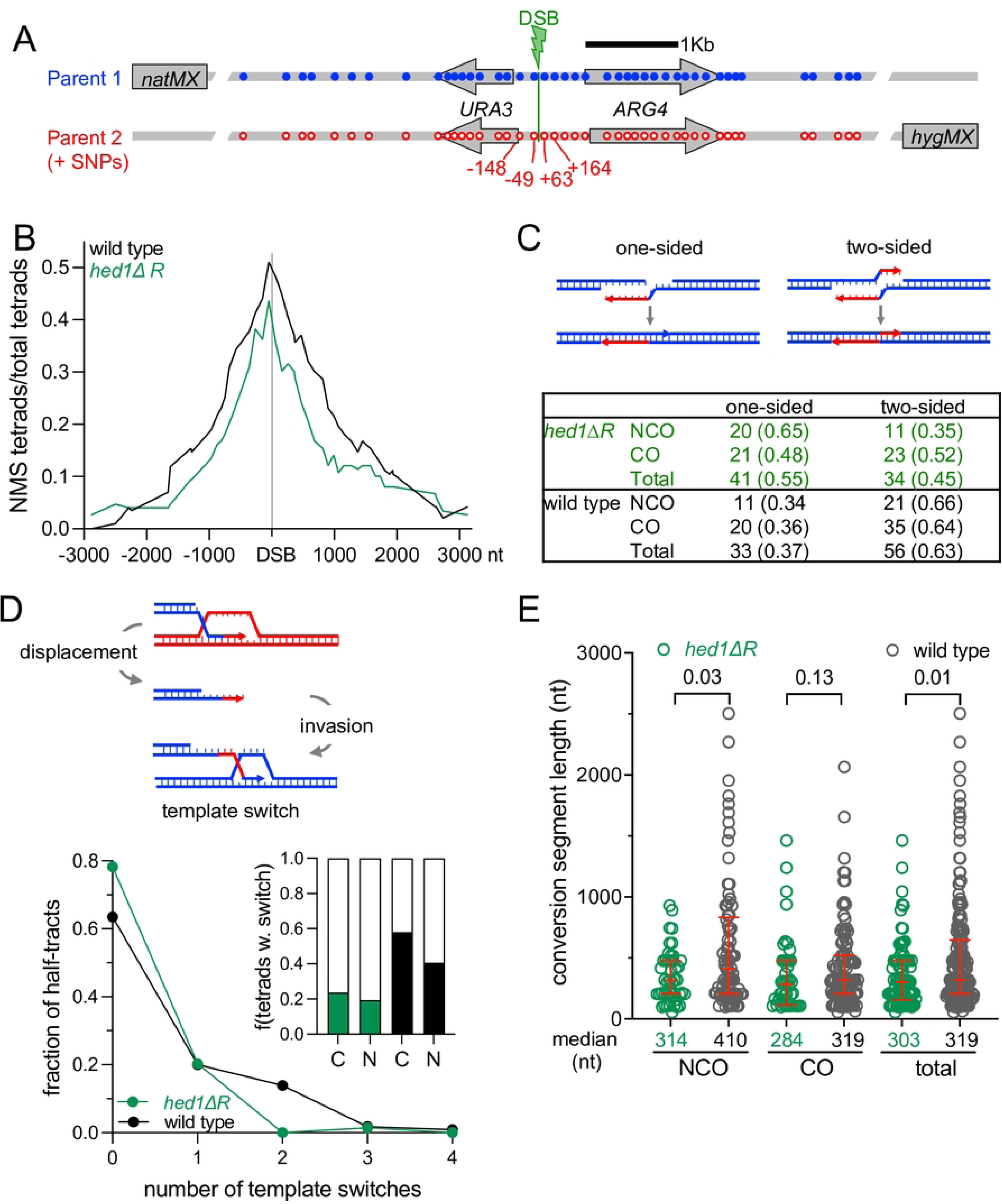
Constitutive activation of Rad51 results in inefficient interhomolog strand invasion during meiosis. (A) The *URA3-tel-ARG4* recombination interval showing the position of polymorphic markers (blue—wild type; red–polymorphisms) relative to the double-strand break centroid (DSB, green lightning bolt). ∼85% of DSBs form between −49 and +63; all DSBs form between −148 and +164 (Ahuja et al., 2021). Flanking drug resistance inserts allow genetic scoring of crossing-over. For the analysis, DNA was isolated from the four spores of 159 tetrads from the *hedΔR msh2Δ* diploid, NH2700. Fragments containing the hotspot region were then amplified and sequenced in their entirety. (B) Gene conversion frequencies (tetrads with nonmendelian segregation (NMS) /total tetrads) for each polymorphic marker. Conversion frequencies in *hedΔR* are significantly less than in wild type (p < 0.0001, Wilcoxon matched-pairs signed rank test). Underlying data are in S2_Data sheet 4 and in [126]. (C) One-sided and two-sided hybrid DNA among noncrossovers (NCO) and crossovers (CO). Schematic—in one-sided events, one DSB end invades a homolog and primes DNA synthesis before disassembly and annealing, leading to hybrid DNA to one side of the DSB; in two-sided events, both ends invade a homolog and extend, leading to hybrid DNA on both sides of the DSB. Table— number of tetrads with one-sided and two-sided events, with fraction of tetrads of each type in parentheses. Individual events are illustrated in Fig S5. More events are one-sided in *hedΔR* than in WT (55% and 37%, respectively; p = 0.02, Fisher’s exact test). Underlying data are in S2_Data, sheet 5 and in [126]. (D) Template switching, expressed as fraction of non-mendelian segregation (NMS) half-tracts (interval from DSB to last converted marker) with the indicated number of template switches. *hedΔR* (green) has fewer template switches than WT (black; p = 0.02, chi-square test). Inset– fraction of tetrads with (filled) or without (clear) template switching (p = 0.003, chi-square test); C—crossovers, N—noncrossovers. Underlying data are in S2_Data, sheets 6 and 7 and in [126]. (E) Length of gene conversion segments, which represent the stretch of DNA between the DSB and the end of a gene conversion tract if no template switching occurs, or, if template switching occurs, the stretch of DNA either between the DSB and a template switch or between two template switches. Median segment lengths are indicated below the X axis; red lines indicate lower, median, and upper quartiles. *p* values (Mann-Whitney test, two-tailed) for comparisons between *hedΔR* (green) and WT (black) are at the top of the plot. Underlying data are in S2_Data, sheet 6 and in [126].

Meiotic recombination frequently involves template switching, where strand invasion switches between homolog and sister chromatids, thus undergoing multiple rounds of invasion, end-primed synthesis, and disassembly [125–130]. Template switching was also reduced in *hedΔR* relative to WT (Fig 7DE, S2_Data, Sheet 7). Comparing *hedΔR* and WT, the average number of template switches per conversion tract (0.25 versus 0.57; *p* = 0.02, chi-square test) and the fraction of events displaying template switching (0.22 versus 0.51; *p* = 0.003, chi-square test) were substantially reduced when Rad51 was constitutively activated. In principle, template switching can be reduced either by decreasing strand invasion or by increasing D-loop disassembly; the latter would be expected to increase the time available for end extension and thus increase the length of gene conversion segments. However, gene conversion segments in *hedΔR* were, if anything, shorter than in WT (Fig 7E, S2_Data, Sheet 6).

In summary, both overall gene conversion frequencies and the fraction of events with multiple strand invasions were reduced in *hedΔR* cells, consistent with the suggestion that meiotic strand invasion is reduced when Rad51 is constitutively activated.

### *PIF1* helicase activity and PCNA interaction promote spore viability and chromosome disjunction when Rad51 is constitutively active

The delay in prophase I observed when Rad51 is constitutively active suggests that processing of Rad51-generated intermediates is different than that of Dmc1-generated D-loops. One possible difference is the use of the Pif1 helicase which plays a key role in extension of Rad51-mediated D-loops in BIR and the formation of a fraction of vegetative crossovers, but has only a minor role in Dmc1-mediated recombination [109, 112]. To see if *PIF1* has a function when Rad51 is constitutively activated during prophase I, a meiotic depletion allele (*pif1-md*) was created by placing *PIF1* under the control of the *CLB2* promoter [112, 131]. *CLB2* is expressed in vegetative cells but not during meiosis [132]. The *pif1-md* mutant alone exhibited ∼85% spore viability, a significant decrease from WT, while Chromosome VIII MI non-disjunction increased only slightly (0.4% vs 0%) (Fig 8AC) [112]. Activation of Rad51 in the absence of *PIF1* (*hedΔR pif1-md*) had no obvious effect on spore viability but significantly increased MI nondisjunction to ∼3%, indicating a synergistic effect (Fig 8AC). Furthermore, depleting *PIF1* in the *hedΔR NDT80-mid* diploid dramatically reduced the average spore viability from ∼60% to ∼25% and increased chromosome VIII MI nondisjunction three-fold compared to *hedΔR NDT80-mid* (Fig 8AC). The effect of *pif1-md* on chromosome segregation was not limited to Chromosome VIII, as the MI non-disjunction pattern of the viable spores in tetrads was exacerbated in the *hedΔR NDT80-mid pif1-md* diploid compared to *hedΔR NDT80-mid* (Fig S6B). Both the spore inviability and MI non-disjunction phenotypes of *hedΔR NDT80-mid pif1-md* were complemented by integration of a *PIF1* plasmid, confirming that depletion of *PIF1* is responsible for these phenotypes (Fig S6AB). In addition, these phenotypes were suppressed by *rad51-II3A*, demonstrating that the more extreme mutant phenotypes observed in the absence of *PIF1* are due to constitutive activation of Rad51 and the absence of the MRC (Fig 8BD).

**Fig 8.**
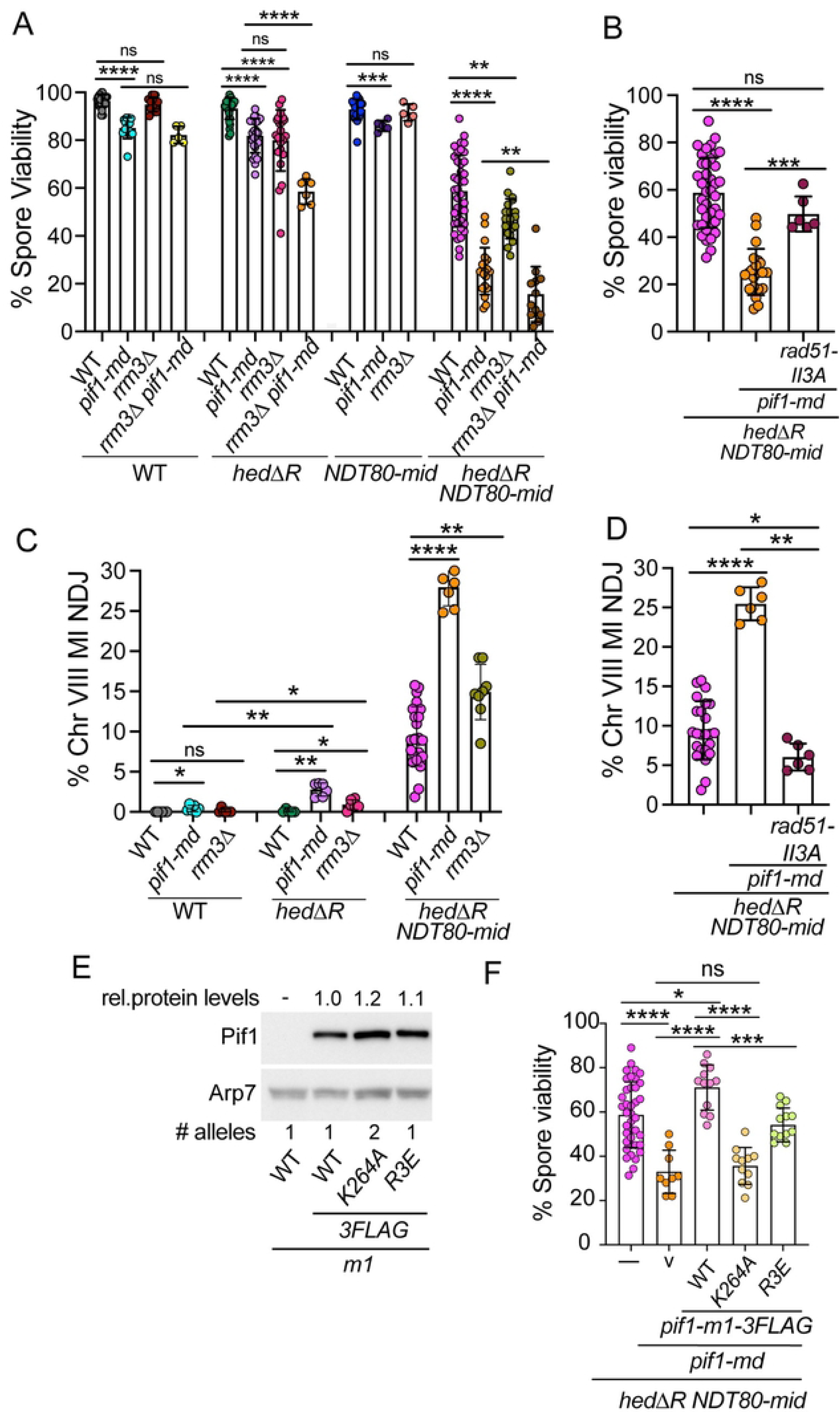
*PIF1* and *RRM3* promote homolog disjunction when Rad51 is constitutively activate in the absence of the meiotic recombination checkpoint. (A) Spore viability. The data from WT, *hedΔR*, *NDT80-mid* and *hedΔR NDT80-mid* were taken from Fig 3E. Additional strains were *pif1-md* (NH2657 or NH2685 RCEN), *rrm3Δ* (NH2485::pRS304^2^ or NH2623 RCEN), *pif1-md rrm3Δ* (NH2671::pRS304^2^), *hedΔR pif1-md* (NH2691 or NH2702 RCEN), *hedΔR rrm3Δ* (NH2540 or NH2629 RCEN), *hedΔR pif1-md rrm3Δ* (NH2704), *NDT80-mid pif1-md* (NH2725), *NDT80-mid rrm3Δ* (NH2549), *hedΔR NDT80-mid pif1-md* (NH2661 or NH2687 RCEN), *hedΔR NDT80-mid rrm3Δ* (NH2596), and *hedΔR NDT80-mid pif1-md rrm3Δ* (NH2670). Sporulated cells from both liquid and solid sporulation media were dissected to determine the percent of viable spores. At least 20 tetrads were dissected for each replicate. (B) Suppression of *hedΔR NDT80-mid pif1-md* spore inviability by *rad51-II3A* (NH2741 RCEN). (C) Frequency of chromosome VIII nondisjunction during MI. The data from WT, *hedΔR*, *NDT80-mid* and *hedΔR NDT80-mid* were taken from Fig 3G. Additional strains used in Panel C were *pif1-md* (NH2685 RCEN), *rrm3Δ* (NH2623 RCEN), *hedΔR pif1-md* (NH2702 RCEN), *hedΔR rrm3Δ* (NH2629 RCEN), *hedΔR NDT80-mid pif1-md* (NH2687 RCEN), and *hedΔR NDT80-mid rrm3Δ* (NH2637 RCEN). Number of tetrads assayed: *pif1-md* (3168), *rrm3Δ* (1062), *hedΔR pif1-md* (1665), *hedΔR rrm3Δ* (900), *hedΔR NDT80-mid pif1-md* (1491) and *hedΔR NDT80-mid rrm3Δ* (1489). (D) Suppression of *hedΔR NDT80-mid pif1-md* Chromosome VIII MI nondisjunction by *rad51-II3A* (NH2741 RCEN). Number of tetrads assay for *hed1ΔR NDT80-mid pif1-md rad51-II3A* (3469). (E) Steady state levels of different nuclear Pif1 proteins from the strains analyzed in Panel F. Immunoblots using protein extracts made from vegetative cells were probed with α- FLAG antibodies to detect Pif1-3FLAG. Arp7 was used as a loading control. A diploid carrying untagged *pif1-m1* (pJW5-m1) was included as a control. The numbers at the bottom of each lane indicate the number of plasmids in each strain. The amount of protein was quantified and normalized to Arp7 and Pif1-m1-3FLAG using the equation [Pif1-m1-X-3FLAG/Arp7]/[Pif1-m1-3FLAG/Arp7], where X indicates a mutation. Numbers above each lane indicate the average values for three independent replicates. (F) Complementation tests for different alleles of *PIF1*. The *hedΔR NDT80-mid pif1-md* diploid was transformed with the vector, pRS306 (v), one copy each of pJW14 (*pif1-m1- 3FLAG*) and pJW14-R3E (*pif1-m1-R3E-3FLAG*), as well as two copies pJW14-K264A (*pif1-m1-K264A-3FLAG*). Transformants were sporulated on solid medium and at least 20 tetrads were dissected per transformant. The statistical significance of differences between strains was determined using the Mann-Whitney test (* = *p*<.05; ** = *p*<.01; *** = *p*<.001, **** = *p*<.0001).

*PIF1* functions both in the nucleus and the mitochondrion [133, 134]. Mitochondrial Pif1 can be eliminated by mutating the first methionine (*pif1-m1*), so that only nuclear Pif1 is present [135]. The *pif1-m1* allele complemented the spore viability defect of *hedΔR NDT80-mid pif1-md* even better than *PIF1*, perhaps because all the Pif1 protein was localized to the nucleus (Fig. S6A). Pif1-m1 was tagged with three FLAG epitopes at the C-terminus (*pif1-m1-3FLAG*) to allow visualization of the nuclear protein. Addition of the tag had no obvious effect on *pif1-m1* function as *pif1-m1* and *pif1-m1-3FLAG* complemented equally well (Fig. S6A).

Pif1 helicase activity and PCNA interaction are both required for BIR [109, 111]. To test whether these functions are similarly required for processing of meiotic Rad51- mediated intermediates, complementation tests were performed in the *hedΔR NDT80- mid pif1-md* diploid using mutant *pif1* alleles that either abolish helicase activity (K264A) or disrupt the Pif1-PCNA interaction (R3E) [109, 111]. Steady state levels of the Pif1- m1-K264A-3FLAG protein were approximately two-fold lower than Pif1-m1-3FLAG and Pif1-m1-R3E-3FLAG (Fig 8E). Complementation tests were therefore performed using diploids hemizygous for the *pif1-m1-3FLAG* WT and *R3E* alleles and homozygous for the *K264A* mutant so that the protein levels of the three proteins were equivalent (Fig 8EF). The *pif1-R3E* mutant only partially complemented. Partial complementation by *pif1-R3E* has been observed previously in other assays and therefore the mutation may not completely disrupt the Pif1-PCNA interaction *in vivo* [111, 112]. Consistent with this idea, recruitment of Pif1 to meiotic DSBs was reduced, but not eliminated, in the *pif1- R3E* mutant [112]. In contrast, the helicase dead *K264A* mutant exhibited the same low spore viability as the vector alone (Fig 8F). We conclude that Pif1 helicase activity and its ability to interact with PCNA are required for processing of Rad51-mediated D-loops during meiotic prophase I.

### *PIF1* promotes meiotic progression and crossover formation in strains with constitutively active Rad51

While similar meiotic progression timing was observed in WT and *pif1-md*, depleting Pif1 from the *hedΔR* background significantly delayed meiotic progression compared to *hedΔR* alone (Fig S7AB). This delay was not due to increased prophase I lifespan (Figure S7C). Even so, eliminating the MRC from the *hedΔR pif1-md* diploid using *NDT80-mid* suppressed the meiotic progression delay to the same T_50_ value as *hedΔR NDT80-mid* and restored prophase lifespan to that of WT (Fig S7). These results suggest activating Rad51 in the absence of *PIF1* may create intermediates that are not repaired before Meiosis I which may then trigger a DNA damage checkpoint delay in Meiosis II [119].

*pif1-md* exhibited WT kinetics for both CO2 and NCO1 formation at the *HIS4LEU2* hotspot, but the levels of both were slightly reduced, perhaps due to the reduced number of cells that entered meiosis (Fig S3, S7A). IH dHJs were decreased, but there was no corresponding increase in IS dHJs (Fig 5C). CO2 formation was delayed in *hedΔR pif1-md* but reached similar levels compared to *hedΔR* and *hedΔR NDT80-mid* (Fig S3). IS dHJs were increased in *hedΔR pif1-md* compared to either *pif1-md* or *hedΔR*, perhaps explaining the small increase in MI non-disjunction observed in this diploid (Fig 5C, 8C). CO2 and NCO1 levels in *hedΔR NDT80-mid pif1-mid* were reduced more than *hed1ΔR NDT80-mid* (Fig S3).

Crossover frequencies were determined using the orthogonal fluorescent spore system was consistent with the physical analysis. The *pif1-md* mutant alone exhibited little to no reduction in map distance in the *CEN8-ARG4* and *ARG4-THR1* intervals, respectively, on chromosome VIII (Fig 4E). In contrast depleting *PIF1* from the *hedΔR* diploid significantly reduced crossovers in the *CEN8-ARG4* interval, and further reductions were observed in both intervals with the *hedΔR NDT80-mid pif1-md* diploid (Fig 4E).

Crossover interference is a genetic phenomenon in which a crossover in one interval lowers the probability of a crossover in an adjacent interval [136]. To analyze interference occurring between crossovers in the *CEN8-ARG4* and *ARG4-THR1* intervals, the method of Malkova, Swanson (137) was used. A crossover interference ratio <1 indicates that interference is occurring. Within each strain the crossover interference ratios calculated using either *CEN8-ARG4* or *ARG4-THR1* as the reference interval were similar (Fig 4F). Crossover interference was observed in WT, *hedΔR*, *NDT80-mid*, *pif1-md*, and *hed1ΔR pif1-md* at levels consistent with the literature (∼0.2) [122]. In contrast, interference was absent in *hedΔR NDT80-mid pif1-md*.

### The *PIF1* paralog, *RRM3*, promotes chromosome segregation when Rad51 is constitutively active during meiosis

*RRM3* and *PIF1* have distinct functions in some processes, e.g., *RRM3* is not required for BIR, and Rrm3 travels with the replisome to remove non-nucleosomal proteins while Pif1 is recruited to the replisome only in specific circumstances [109, 138–141]. However, there are also examples where *PIF1* and *RRM3* appear to function redundantly [142, 143]. A role for *RRM3* in meiosis was therefore examined. No effect on spore viability was observed in *rrm3Δ* or *NDT80-mid rrm3Δ* cells (Fig 8A). In contrast, a small, but significant, decrease in spore viability was observed in *hedΔR rrm3Δ* (Fig 8A). In addition, there was a corresponding increase in Chromosome 8 MI non-disjunction (Fig 8C). These phenotypes were exacerbated when the meiotic recombination checkpoint was removed (*hedΔR NDT80-mid rrm3Δ*) (Fig 8AC, S6C). Both the spore inviability and MI nondisjunction phenotypes were complemented by addition of *RRM3* on a plasmid (Fig S6AC). The spore viability defect was not the result of mutations that accumulated during vegetative growth in the absence of *RRM3*, since the spore inviability phenotype of the *hedΔR NDT80-mid rrm3Δ* was complemented by expressing *RRM3* specifically during meiosis using the *REC8* promoter (Fig S6A).

If *PIF1* and *RRM3* function redundantly in processing Rad51 generated intermediates during meiosis, a synergistic defect in spore viability when the two mutants were combined is expected. Alternatively, if they function independently of each other, the predicted spore viability of *pif1-md rrm3Δ* in various backgrounds would be the product of their individual spore viabilities. A diploid containing *pif1-md* and *rrm3Δ* exhibited the same level of viable spores as *pif1-md* alone, consistent with the lack of any phenotype for *rrm3Δ* in an otherwise WT diploid (Fig 8A). In contrast, in the *hedΔR* and *hedΔR NDT80-mid* backgrounds, the average spore viabilities for the *pif1-md rrm3Δ* containing diploids were close to the product of the individual values (*hedΔR pif1-md* x *hedΔR rrm3Δ*, expected: 82% * 80% = 66%; observed for *hedΔR pif1-md rrm3Δ* 59%; *hedΔR NDT80-mid*, expected: 25% * 47% = 12%; *hedΔR NDT80-mid pif1-md rrm3Δ* observed 16%). We conclude that *PIF1* and *RRM3* play different roles in processing Rad51-mediated intermediates.

## Discussion

### The meiotic recombination checkpoint promotes proper chromosome segregation and spore viability in WT and *hedΔR* diploids

It has often been suggested that the MRC functions to delay prophase I exit to allow time for DSB repair to occur. One difficulty in proving this idea, however, has been the pleiotropic nature of the Mek1 kinase. *MEK1* is required not only to regulate Ndt80 and therefore prophase I exit, but also for interhomolog bias [75]. Inactivation of Mek1, through mutation or kinase inhibition, results in rapid repair of DSBs via the sister chromatids, thereby removing the signal to the checkpoint and allowing efficient meiotic progression [77–79]. In addition, mutants that disrupt the MRC such as *RAD24*, *RAD17*, and *MEC1/TEL1* all play a role in Mek1 activation and therefore their effects could be indirect due to affecting other *MEK1* processes as well [92, 144–146]. Artificially prolonging the length of prophase I using an inducible allele of *NDT80* was found to improve the spore viability of *mec1* and *rad24* mutants, and this condition was suggested to mimic the prophase I delay caused by the meiotic recombination checkpoint [147]. A problem with this interpretation, however, is that cells behave aberrantly during the extended prophase I arrest that occurs when *NDT80* is inactive. For example, in a *mek1Δ ndt80Δ* diploid, intersister joint molecules predominate early in prophase I but this bias is switched to interhomolog JMs with increasing time in the arrested cells [148]. Consistent with this result, *mek1Δ* spore inviability can be rescued by allowing increased time in prophase I prior to *NDT80* induction [94]. Our work instead utilized the *NDT80-mid* mutant, which abolishes the MRC without affecting Mek1 kinase activity, to examine the role the MRC plays in WT cells, as well as the *hedΔR* mutant that exhibits a modest defect in DSB repair where meiotic progression is delayed but not arrested.

Prophase I length in the *NDT80-mid* mutant was not significantly shorter than WT. However, spore viability was decreased by a small, but significant amount, as were crossovers in the *ARG4-THR1* interval, and MI non-disjunction of Chromosome VIII was slightly elevated. These results indicate that there are DSB repair problems in a small number of otherwise WT cells that require time provided by the MRC to resolve. Unexpectedly, cultures derived from different single colonies from a WT diploid displayed variability in the length of prophase I ranging from 2-5 hours (Fig 6A). However, high levels of viable spores were observed in all cultures, whether they went fast or slow, suggesting that some cultures completed recombination earlier and exited prophase I, while others needed more time for DSB repair. Variability was also observed in *hedΔR*, although in this case prophase I length was longer, ranging from 3-6 hours. Nevertheless, spore viability remained high (Fig 6A). In contrast, there was a striking, statistically significant, positive correlation between prophase I lifespan and spore viability and a negative correlation between prophase I lifespan and MI non-disjunction when the meiotic recombination checkpoint was abolished in *hedΔR NDT80-mid* (Fig 6AB). These correlations were eliminated in *hedΔR rad51-II3A NDT80-mid* consistent with the fact that Rad51 strand exchange activity was responsible for the reduced spore viability. These experiments demonstrate that a key function of the MRC is to provide time for processing recombination intermediates during meiotic DSB repair.

### Interaction between Rad51 and Rad54 in Phase I allows Rad51-mediated interhomolog strand invasion in the presence of Dmc1

Two major differences between mitotic and meiotic recombination in yeast are that the presynaptic filament contains patches of both Rad51 and Dmc1 and strand invasion is biased toward the homolog and not sister chromatids. The interplay between these recombinases is complex, however: while the presence of the Rad51 protein is required for proper Dmc1 loading and interhomolog bias, Rad51 strand exchange activity is not required for interhomolog recombination and is actively suppressed [25, 30, 84, 86, 87]. In addition, although deletion of *DMC1* abolishes interhomolog strand invasion, full activation of Rad51 in a *dmc1Δ* mutant results in significantly higher levels of interhomolog recombination than *mek1Δ*, indicating that activated Rad51 can generate interhomolog intermediates when Mek1 is active [88, 89]. The modest phenotypes exhibited by *hed1Δ* and *hedΔR* diploids have been attributed to the presence of Dmc1 itself contributing to the inhibition of Rad51 strand exchange activity [88, 89]. However, the idea that activating Rad51 should decrease interhomolog recombination in the presence of Dmc1 assumes that Rad51 would preferentially direct strand invasion to sister chromatids, as it does in vegetative cells. This work shows that Rad51 strand exchange activity is not suppressed by Dmc1 in yeast. Instead, the near WT levels of crossovers and spore viability exhibited by *hed1Δ* and *hedΔR* diploids are due to a combination of different factors that promote the creation of Rad51-mediated interhomolog crossovers during Phase 1.

The fact that specific elimination of the MRC in a *hedΔR* diploid decreased interhomolog events, increased MI non-disjunction and decreased spore viability indicates that Rad51 is generating intermediates that the MRC is providing time to process. Proof of this idea is that all of these mutant phenotypes were suppressed by preventing Rad51 exchange activity using the *rad51-II3A* mutant. Therefore, in these cells, Rad51 is active and generating crossovers during Phase 1 when interhomolog bias is mediated by Mek1. We suggest that the high levels of crossovers and spore viability observed in *hed1Δ* diploids compared to *hed1Δ dmc1Δ* result from the presence of Dmc1 helping the presynaptic filament search for homology on homologs rather than sister chromatids through accessory factors like Hop2/Mnd1 [149].

There are some organisms such as the nematode, *Caenorhabditis elegans*, the fruit fly, *Drosophila melanogaster*, and the fungus, *Sordaria macrospora*, that use only Rad51 for meiotic recombination [3, 150]. This work raises the interesting question of whether Rad51 mediated meiotic recombination in these organisms occurs similarly to the pathway we have discovered in yeast. This idea could be tested by determining whether Pif1 orthologs are required for meiotic recombination in these species.

### A model for Rad51-mediated recombination during meiosis

We propose that processing of nascent meiotic D-loops generated by Rad51 is different from D-loops made by Dmc1 for several reasons. First, activation of Rad51 creates a delay in prophase I exit, indicating that the cell requires more time to satisfy the MRC when there is Rad51-strand exchange activity than when Dmc1 is the sole functioning recombinase. Second, the MRC plays only a minor role in promoting chromosome disjunction and spore viability in WT cells when interhomolog recombination is mediated by Dmc1 but is very important when Rad51 also is active during Phase 1. Third, *PIF1* activity is normally inhibited such that its contribution to Dmc1-mediated repair is negligible [112].

Our model is based on the assumption that similar to mice, Rad51 and Dmc1 form adjacent patches in the presynaptic filament with Dmc1 closest to the 3’ end where it is responsible for strand invasion (Fig 9A) [19, 22]. After the homology search results in formation of a nascent D-loop (Fig 9B), Dmc1 is removed from the 3’ end by a DNA translocase to form heteroduplex between the invading strand and the complementary donor strand, resulting in the displacement of the strand of like polarity to make a mature D-loop (Fig 9C). It is unclear what DNA translocase is responsible for this step. While Rdh54 seems the obvious choice given *in vivo* and *in vitro* evidence that it preferentially works with Dmc1, this activity has not yet been demonstrated for Rdh54 [32, 151]. The 3’ end of the invading strand then acts as a primer for DNA synthesis using PCNA and Polδ to extend the invading strand (Fig 9D) [52–55]. Disassembly of the D-loop followed by annealing to the other side of the break creates noncrossovers, while second end capture of the extended end generates double Holliday junctions that can be resolved as crossovers (Fig 9E). Note that our model shows a simplified view of the recombination process. Recent work has demonstrated that meiotic DSB repair is very dynamic, with multiple rounds of strand invasion and disassembly, and template switching between homolog and sister chromatids [125–127].

**Fig 9.**
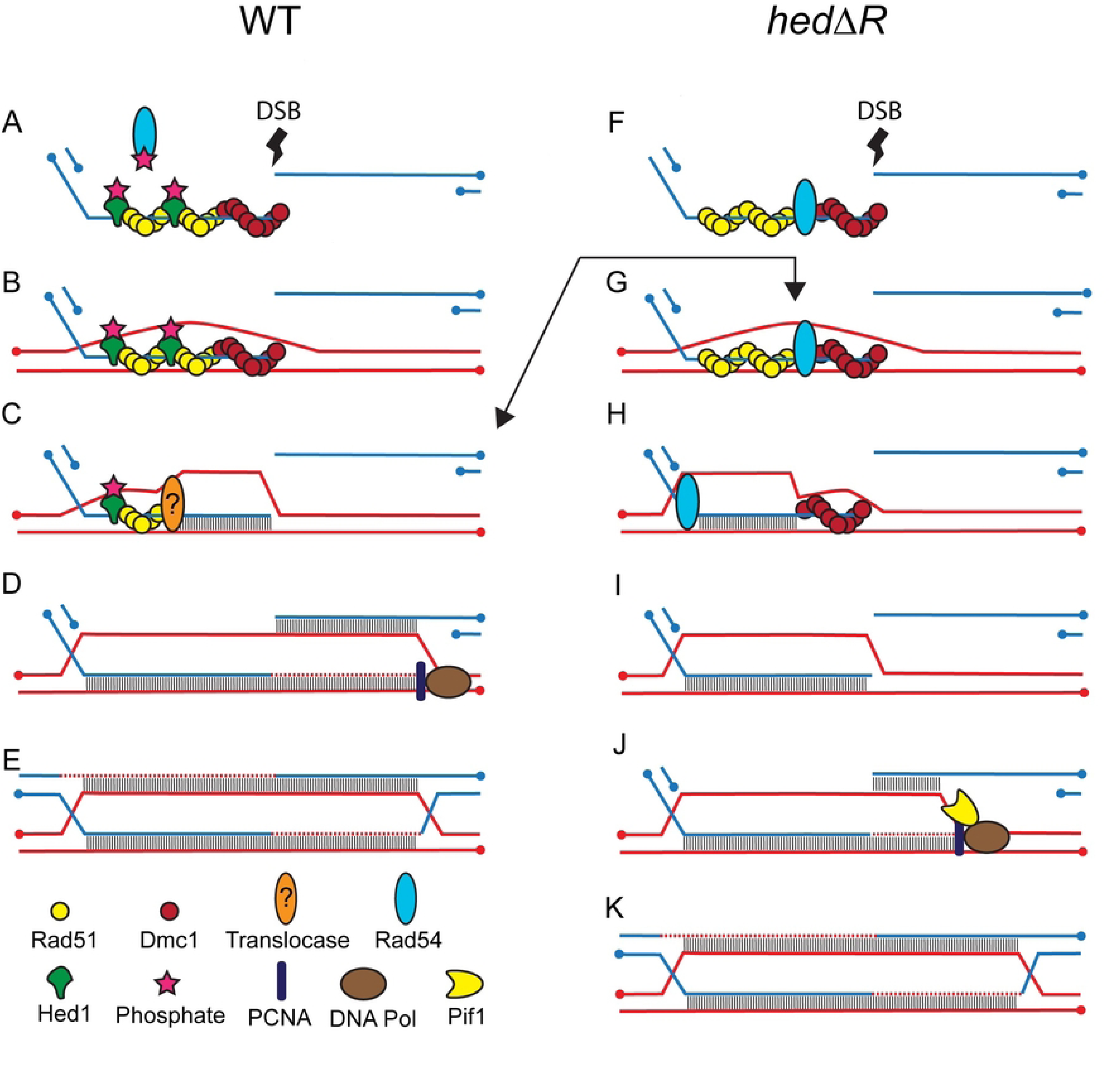
Model for Rad51-mediated interhomolog crossovers during meiosis. (A) Presynaptic filament shown for one side of a DSB in WT cells. Blue lines represent the duplex of one homolog while red lines represent a chromatid from the other homolog. Dots indicate the 5’ ends of the DNA. (B) Nascent D-loop or paranemic joint. (C) Conversion to a mature D-loop by removal of Dmc1 and heteroduplex formation between the invading strand and donor strand of opposite polarity. The question mark indicates that the mechanism for how this occurs is unknown. (D) Extension of the D-loop by DNA synthesis by Polδ-PCNA followed by second end capture. (E) Double Holliday junction. (F) Presynaptic filament shown for one side of a DSB in *hedΔR* cells with Rad54 interacting with Rad51. (G) Nascent D-loop or paranemic joint. (H) Internal D-loop created by Rad54 removal of Rad51 and heteroduplex formation between the invading strand and donor strand of opposite polarity. (I) Removal of Dmc1 from the 3’ end of the invading strand. (J) Pif1 interacts with PCNA to facilitate DNA synthesis by unwinding the duplex to extend the D-loop allowing second end capture. (K) Double Holliday junction. Not shown is the possibility that the extended ends could be disassembled to enable noncrossover formation by synthesis dependent strand annealing.

Interaction between Rad54 and Rad51 is critical for Rad51-mediated repair in meiosis during Phase 2 (Prugar et al., 2017). Rad54 binds to gaps between patches of Rad51 on the ssDNA In vitro [118, 152]. We propose that Rad54 similarly binds in gaps present between Rad51 and Dmc1 on the presynaptic filament (Fig 9F). The resulting presynaptic filaments then form nascent D-loops similarly to WT (Fig 9BG). However, in a subset of DSBs, Rad54 sequentially removes Rad51 monomers, thereby creating heteroduplex DNA between the invading strand and donor strand of opposite polarity to generate an internal D-loop (Fig 9H). The formation of internal D-loops by Rad51-Rad54 has previously been demonstrated *in vitro* [37].

Formation of an internal D-loop by Rad51 creates a problem in that the 3’ end remains bound by Dmc1, thereby preventing it from serving as a primer for DNA synthesis by DNA Polδ (Fig 9H). In addition, the absence of a free 3’ end may make this intermediate more susceptible to disassembly by the STR complex which would explain why there are fewer two-sided events and template switches in *hedΔR*. Having to remove the Dmc1 to free up the 3’ end (Fig 9I) is an extra step that would delay strand extension, thereby taking longer for second end capture to occur (Fig 9J) which could explain the delay in dHJ formation observed for *hedΔR*. In addition, the increased time it takes to convert the break into an intermediate that is not recognized by the MRC would result in an MRC-dependent prophase I delay.

Pif1 is known to move in the 5’-3’ direction to remove proteins from DNA, so its function in processing Rad51-generated intermediates could be the removal of Dmc1 after internal D-loop formation. However, this activity was not detected for Pif1 using *in vitro* experiments with DNA curtains comprised of Dmc1-bound ssDNA, making this possibility less likely (Eric Greene, personal communication). Another possibility is that the end containing Dmc1 is cleaved off to create a free 3’ end located upstream of the DSB site. Extension of this end is predicted to produce 3:1 gene conversions around the DSB site which was not observed. The simplest explanation is that Dmc1 is removed by the same mechanism shown in Fig 9C.

Pif1 activity is normally inhibited during Dmc1-mediated recombination by Mer3 and MutLβ [112]. It is possible that these proteins do not recognize a Rad51-generated internal D-loop and therefore are not present to inhibit Pif1, allowing Pif1 to promote synthesis of the invading strand by binding to PCNA and unwinding the DNA (Fig 9J). The resulting extended strand could then be processed in a similar way used for Dmc1-generated D-loops to make crossovers and noncrossovers (Fig 9K). A key test of the model will be to see if conversion track length is decreased in a *hedΔR pif1-md* diploid.

### Is downregulation of Rad51 activity conserved in mammals?

No ortholog of Mek1 has been reported in mammals, raising the question of whether Rad51 activity is inhibited during prophase I in mammalian cells. Several observations suggest that the answer is yes. First of all, repair of exogenous DNA damage in mouse oocytes exhibits two temporally distinct processes during pachytene—an early one that occurs in the presence of both Dmc1 and Rad51 and a later stage where Rad51, but not Dmc1, is associated with the DSBs [106]. In addition, depletion of Rad51 specifically in mouse spermatocytes results in apoptosis early in prophase I [153]. This phenotype is very different from the zygotene arrest exhibited by Dmc1 depletion or *Dmc1^-/^* spermatocytes [14, 153, 154]. These results show that (1) the presence of Rad51 is required for Dmc1-mediated recombination and (2) that Rad51 is not mediating strand invasion when Dmc1 is absent during Phase I, suggesting that its strand exchange is being inhibited similar to yeast. The mechanism of this inhibition is unclear, but it requires a functional chromosome axis as mutants in genes encoding the mouse axial element proteins Sycp2 and Sycp3 allow *Rad54*^+/+^-dependent intersister repair of DSBs in *Dmc1^-/-^* oocytes [155].

In summary, Rad51 activity is tightly regulated during meiosis to prevent it from competing with Dmc1 in the repair of DSBs during Phase 1, while allowing Rad51 activity at the end of prophase I in Phase 2 to rapidly repair any remaining breaks prior to MI. These two phases of DSB repair are coordinated via the MRC through Mek1 regulation of the Ndt80 transcription factor. Rad51-mediated repair in meiotic cells has similarities with BIR in vegetative cells in the use of the Pif1 helicase. Pif1 is conserved in mammals where it is also required for BIR [156]. It would be interesting to see if it has a role in the repair of DSBs in *Dmc1^-/-^ Sycp2^-/-^* mouse oocytes.

## Materials and Methods

### Strains

All strains were derived from the SK1 background and their genotypes are listed in Table S1. Liquid and solid media are described in [157]. Polymerase chain reaction (PCR) mediated gene deletions were generated using the drug resistance markers *kanMX6, natMX4* and *hphMX4* which confer resistance to the antibiotics G418, nourseothricin (NAT) and Hygromicin B (HygB), respectively [158, 159]. Deletions were confirmed by PCR using a forward primer upstream of the gene’s open reading frame (ORF) and a reverse primer located within the drug marker, while the absence of the WT allele was assayed using the same forward primer and a reverse primer located within the open reading frame (ORF). The *pif1-md* allele was constructed by fusing the WT *PIF1* ORF to the *CLB2* promoter. A *kanMX6*-*P_CLB2_-3HA* PCR fragment with 50 bp homology from −100 to −50 from the *PIF1* ATG on the upstream end and 50 bp homology from +1 to +50 from the ATG on the downstream end was amplified using pMJ787 (also called 455) [131] and transformed into haploid yeast strains selecting for G418 resistance. The gene fusion was confirmed by colony PCR using a forward primer in the *CLB2* promoter and a reverse primer in the *PIF1* ORF.

The *NDT80-mid*, *RAD54-T132A* and *rad51-II3A* alleles were introduced into haploid strains by two-step gene replacement [160]. The *NDT80-mid URA3* integrating plasmid pNH317 was digested with SnaBI to target integration of the plasmid upstream of *ndt80Δ::kanMX6*. Ura^+^ transformants were grown in YPD and then plated on 5-fluoro-orotic acid (5-FOA) to select for cells that had lost the *URA3* plasmid [161]. Foa^R^ colonies that retained the *NDT80-mid* allele were identified by loss of the *ndt80Δ::kanMX6* marker, rendering the cells sensitive to G418.

The *RAD54-T132A URA3* integrating plasmid, pHN104, was digested with BsiWI to target integration upstream of *RAD54*. Ura^+^ transformants were grown in YPD and then plated on 5-FOA. A fragment of the *RAD54* gene from each Foa^R^ colony was amplified by PCR and sequenced to detect the presence of the T132A mutation. With the exception of the PacBio sequencing used for the DSB hotspot analysis (see below), all DNA sequencing was performed by the Stony Brook University DNA Sequencing Facility.

The *rad51-II3A URA3* integrating plasmid, pAM1-II3A, was digested with SpeI to target integration upstream of *RAD51*. Ura^+^ transformants were grown in YPD then plated on 5-FOA. Foa^R^ colonies were patched onto YPD and replica plated to YPD+0.04% methyl methanesulfonate (MMS) to screen for MMS sensitivity. The *rad51- II3A* allele contains three mutations, R188A, K361A and K371A. To confirm that the Mms^S^ Foa^R^ cells contained all three mutations, the *RAD51* gene was amplified from genomic DNA and sequenced.

All of the strains used for fluorescent spore analysis are isogenic with NH716, which is created by crossing the NHY1210 and NYH1215 haploids. To allow integration of the appropriate plasmids, the *LEU2* and *TRP1* genes were deleted from NYH1210 and NHY1215 (and their mutant derivatives), respectively. All but the first and last 100 bp of the *LEU2* ORF ectopically located in the NHY1210 haploid were replaced with *kanMXM6* (designated *leu2ΔI::kanMX6*). Additionally, the 5’ end of the *TRP1* gene was replaced with *hphMX4* in the NHY1215 haploid [80].

To measure the frequency of chromosome VIII nondisjunction [122], the plasmids, pSK693 (*pYKL050c-RFP LEU2*) and pSK694 (*pYKL050c-CFP TRP1*) were digested with AflII to target integration to the right arm of chromosome VIII adjacent to the centromere in NHY1210 and NYH1215 (and their mutant derivatives), respectively. Diploids with the *RFP* and *CFP* genes in allelic positions are indicated by “RCEN”. To measure genetic map distances between *CEN8-ARG4* and *ARG4-*THR1, pSK729 (*pYKL050c-GFP* URA3*) and pSK695 (*pYKL050c-CFP TRP1*) were digested with AflII and sequentially integrated into NHY1215 haploids at *ARG4* and *THR1*, respectively. These haploids were crossed to the appropriate NHY1210 CEN8::RFP haploids to create diploids containing the marker configuration indicated by “RGC”.

The *hedΔR msh2Δ* haploids used for the DSB hotspot experiment were created in several steps. First the *HED1* gene was deleted from S5495 and S5497 using *natMX4*. To inactivate *natMX4* so this gene could be used in a later step, the *hed1Δ::natMX4* haploids were then transformed with pSZ5, a 2μ plasmid containing *LEU2* and *CAS9* fused to 5’TTCGTCGCGTACGGGGACGA3’, a guide DNA sequence that, once transcribed, targets DSB formation within *natMX4*. Because the broken chromosome is lethal, there is a strong selection for cells in which error-prone non-homologous end joining has occurred that destroys the Cas9 cut site and disrupts the *natMX4* ORF. Leu^+^ transformants were patched onto SD-leu plates and then replica-plated to YPD + NAT to look for patches that failed to grow. Nat^S^ (NS) patches were streaked for single colonies on YPD that were then replica plated to SD-leu to find cells that had lost pSZ5. The *RAD54-T132A* allele was introduced into S5495 hed1NS and S5497 hed1NS as described above. The resulting strains were designated S5495 h*Δ*R and S5495 h*Δ*R. The *hed1Δ::NS RAD54-T132A* mutants were then introduced into haploid strains containing *msh2Δ* and the *URA3-tel-ARG4* hotspot with or without SNPs. First, S5495 h*Δ*R was crossed to S4955 to make NH2694. This diploid was heterozygous for *hed1Δ::NS*, *RAD54-T132A*, as well as *leu2-R*, *his4Δ::kanMX*, *his4*::*URA3rev-tel1-ARG4 +SNPs-RNQ-hphMX-FUS1* and *msh2Δ:kanMX6*. NH2694 was sporulated and 100 tetrads dissected. Strains that were Ura^+^, Arg^+^, HygB^R^ and G418^R^ were then screened for mating type and *MAT***a** strains were selected. (The *his4Δ::kanMX* gene is on chromosome III, as is *his4*::*URA3rev-tel1-ARG4 +SNPs-RNQ- hphMX-FUS1*, so it should not be present in this group). The presence of *msh2Δ::kanMX6* and *hed1Δ::NS* was confirmed by PCR, while spore colonies containing *RAD54-T132A* were detected by DNA sequencing. The resulting haploid, NH2694-a2 was frozen down. To construct the other parent, S5497 h*Δ*R was crossed to S5085 to generate NH2685. This diploid was heterozygous for *hed1Δ::NS RAD54- T132A*, as well as *his4Δ::kanMX*, *STE50-natMX-RRP7*-*his4*::*URA3rev-tel1-ARG4* and *msh2Δ:kanMX6*. 100 tetrads from NH2685 were dissected and the spore colonies screened for Nat^R^ Ura^+^ Arg^+^ and G418^R^. From this group, *MATα* strains were selected and tested for *msh2Δ::kanMX6* and *hed1Δ::NS* by PCR. The presence of *RAD54- T132A* was determined by DNA sequencing. The haploid NH2695-f8 was frozen down.

Prior to transformation, plasmids containing *PIF1* in the *URA3* integrating vector, pRS306, were digested with BmgBI to target integration downstream of the *P_CLB2_-PIF1* (*pif1-md*) allele. Similarly, *URA3* integrating plasmids containing various alleles of *RRM3* were digested with SnaBI to target integration downstream of *rrm3Δ*. The presence of the introduced allele was determined by PCR using a locus-specific primer and a gene-specific primer. For the vector control, pRS306 was digested with NsiI and integrated downstream of *ura3*.

### Plasmids

The genotypes of the plasmids used in this study are listed in S2_Table. The pAM1-II3A plasmid containing the *rad51-II3A* (*rad51-R188A-K361A-K371A*) allele from Cloud, Chan (25) was created by PCR site-directed mutagenesis. First a 3.7 kb BamHI fragment containing *RAD51* was cloned into the YIp5 plasmid backbone digested with BamHI to create pAM1. Site-directed mutagenesis was used on pAM1 to subsequently introduce R188A, then K621A and finally K371A to create pAM1-II3A. All mutations introduced into pAM1-II3A were confirmed by DNA sequencing.

The pNH257 plasmid contains the *REC8* promoter in a 266 bp fragment located immediately upstream of the *REC8* ORF. This plasmid was created by amplifying this fragment from a plasmid containing *REC8* with Not1 and BamHI restriction sites engineered on the ends. The BamHI site is located immediately upstream of where the *REC8* start codon would be. This fragment was cloned into NotI/BamHI digested pRS306.

All of the plasmids created by Gibson assembly used the NEBuilder HiFi DNA Assembly kit (New England Biolabs, #E2621). To construct the *URA3 RRM3* plasmid, pBG22, a 3.5 kb fragment containing the *RRM3* ORF and 900 bp 5’ flanking sequence and 400 bp of 3’ flanking sequence was amplified from pBG18 and cloned by DNA assembly into pRS306 digested with EcoRI and ClaI. pJW7 was constructed by cloning a fragment containing the *RRM3* ORF and 378 bp of downstream sequence 18 bp downstream of the *REC8* promoter in pNH257 vector digested with EcoRI and ClaI using the Gibson Assembly method.

pJW5 was constructed by first amplifying the entire *PIF1* gene with 904 bp and 401 bp of upstream and downstream flanking sequences, respectively, from SK1 genomic DNA. This fragment was cloned into pRS306 digested with EcoRI and XhoI by Gibson Assembly. Mutant derivatives of pJW5 were also generated by Gibson Assembly by introducing two fragments that partitioned the insert employed for the pJW5 construction: one stretching from 904 bp upstream of the *PIF1* open reading frame (ORF) to the mutation and another stretching from the mutation to 401 bp downstream of the *PIF1* ORF, into pRS306 digested with EcoRI and XhoI. pJW14 (*pif1- m1-3FLAG*) was constructed by Gibson Assembly cloning of three fragments into EcoRI-HindIII-digested pRS306: fragment 1 stretches from 904 bp upstream of the *PIF1* ORF to the *pif1-m1* mutation (M1A). Fragment 2 stretches from the *pif1-m1* mutation to the *3xFLAG* tag sequence and fragment 3 stretches from the *3xFLAG* tag sequence to 401 bp downstream of the *PIF1* ORF. Mutant derivatives of pJW14 were constructed by a similar workflow to pJW14 except that Fragment 2 was amplified from derivatives of pJW5 containing either the K264A (AAA to GCG) or R3E (I817R, M820R, L821R, R823E). pJW11 (*URA3 PIF1-3FLAG*) was constructed by amplifying two fragments, one of which contains the *PIF1* gene and the other the 3FLAG sequence and cloning them by Gibson assembly into pRS306 digested with EcoRI and XhoI. All plasmid constructions were confirmed by DNA sequencing at the Stony Brook University DNA sequencing facility.

### Yeast media and meiotic timecourses

Yeast media and sporulation protocol are described in [157]. YPDcom plates contain 2% agar, 0.7% yeast nitrogen base without amino acids, 2% glucose and 2 g complete powder made as described in [157].

### Immunoblots and antibodies

Protein extracts were generated using the trichloroacetic acid method described in [162]. The Rad51 antibody was generated by Covance Research Products (now Labcorp) in guinea pigs using the peptide QKDGNDFKFNYQGDEC. The specificity of the antibody for immunofluorescence of chromosome spreads was demonstrated by the lack of foci observed at the 4.5 hr meiotic timepoint in a *rad51Δ* diploid, compared to numerous foci that were present at the same timepoint in WT (Fig S1A). For the Pif1- m1-3FLAG immunoblots, mouse α-FLAG antibodies (Sigma F-1804) were used at a 1:5000 dilution at room temperature for two hours. The secondary antibody was goat α-mouse (Invitrogen, #PI32430), used at 1:10,000 dilution for one hour at room temperature. Arp7 protein was used a loading control and detected using α-Arp7 primary antibodies (Santa Cruz Biotechnology, #SC8961) with mouse α-goat secondary antibodies (Santa Cruz Biotechnology, #SC2354) used at 1:10,000 dilution at room temperature for 1 hr. Proteins were detected by a chemiluminescent assay using the Advansta Westernbright ECL kit from Fisher (#K-12045-D20).

### Measuring meiotic progression and whole cell immunofluorescence

1.5 ml of 4.5×10^7^ sporulating cells from meiotic timecourses were fixed overnight at 4°C with a final concentration of 3.7% formaldehyde at various timepoints. The next day, fixed cells were washed once with 1 ml phosphate buffered saline (PBS) and stored in 1.5 ml PBS at 4°C. For analysis of meiotic progression using DAPI, 30 µl of fixed cells were placed onto each well of coated slides (Carlson Scientific cat# 101206). Cells were allowed to settle onto the slides for 15 min at room temperature (RT). After incubation, the remaining liquid was aspirated away, and slides were air dried. 3 µl of DAPI plus mounting medium (Vectashield H-1200) was added to each well and the slide was covered with a coverslip and sealed with nail polish. DAPI stained cells were observed using a Zeiss Axio Imager.Z2 microscope with a Zeiss Plan-Apochromat 63X objective. 200 cells were counted at each timepoint and categorized as either mononucleate, binucleate (meiosis I) or tetranucleate (meiosis II). T_50_ values, or the amount of time in hr for a particular sporulation culture to reach ½ its maximum %MI+MII value, were determined graphically for each culture.

For whole cell immunofluorescence analysis, coated slides containing 12 wells (Carlson Scientific cat# 101206) were treated by pipetting 30 μl 0.01% poly-L-lysine (Sigma Cat# p4707) into each well and incubating at RT for 15 min. Residual poly-L- lysine was removed from the wells using an aspirator. The slide was then washed twice by pipetting 30 µl of water into each well, followed by aspiration.

For spheroplasting, 1.5×10^7^ fixed cells were incubated in 75 µl ZK buffer (25 mM Tris-Cl pH7.5, 0.8 M KCl) + 0.04 M Dithiothreitol (DTT) for 2 min at room temperature. The cells were pelleted, and the supernatant was removed. The cell pellets were resuspended in 75 µl ZK buffer + 0.1 mg/mL zymolyase 100T (US Biological Z1004) and incubated at room temperature (RT) for 10 min. Spheroplasted cells were centrifuged at low speed (900 x g) and washed 1 X in 500 μl ice-cold sorbitol/MES buffer (1M Sorbitol, 0.1M MES pH6.5, 1 mM EDTA, 0.5 mM MgCl_2_). After washing, cells were resuspended in 150 μl sorbitol/MES + 0.1% Triton X-100 and incubated 5 min at room temperature to permeabilize the cell membrane. The sorbitol/MES buffer wash was repeated.

Permeabilized cells were resuspended in 40 µl sorbitol/MES buffer. 30 µl of resuspended cells was added onto each well of a printed microscope slide. After allowing the cells to settle onto the slide for 15 min, the remaining liquid was aspirated away. The slides were placed into a coplin jar with methanol at −20°C for 6 min and then transferred to a coplin jar with acetone at −20°C for 30 seconds. Afterwards, slides were air dried until all the residual acetone evaporated. 30 µl of 1% Bovine serum albumin (BSA) in Tris buffered saline (TBS) (140 mM NaCl, 2.7 mM KCl, 25 mM Tris pH8.0) was added to each well and incubated for 30 min at room temperature in a slide box with moist paper towels. After incubation, slides were washed in TBS buffer for 30 sec. Excess TBS was removed by leaning the slides on an edge with a paper towel. The rabbit α-Red1 polyclonal antibody (rabbit 16440) (Wan et al. 2004) was diluted 1:300 in 1% BSA. 80 µl of primary antibody dilution was added to the entire slide, then a 25×50 mm coverslip was placed onto the slide. Primary antibodies were incubated overnight (∼14-18 hrs) at 4°C in slide box with moist paper towels. The next day, coverslips were removed in TBS and slides washed 2 X in TBS for 10 min. The goat α-rabbit Alexa 488 antibody (Fisher Cat# A11008) was diluted 1:1000 in 1% BSA. 80 µl of secondary antibody dilution was added to each slide and covered with a coverslip. Secondary antibodies were incubated for 2 hr at 4°C in slide box with moist paper towels. After the secondary antibody incubation, coverslips were removed in TBS and slides were washed 2X in TBS for 10 min. Slides were completely air dried in the dark and 3 µL DAPI plus mounting medium was added to each well. A coverslip was added to the slide and sealed with nail polish. Microscopy images were taken using a Zeiss Axio Imager.Z2 microscope with a Zeiss Plan-Apochromat 63X objective.

To analyze the length of prophase I in different timecourses, 200 cells were counted at each timepoint and classified as either being Red1^+^ or Red1^-^. The %-Red1^+^ cells were graphed over time and the area under the curve (AUC) was calculated using GraphPad Prism software. The AUC for each timecourse was divided by the maximum %MI+MII value to normalize for the number of cells that entered meiosis for that timecourse. This calculation yields the “prophase I lifespan” (hr) and is analogous to the method previously used to calculate the length of pre-meiotic S phase [113].

### Nuclear spreads

Nuclear spreads were prepared as described in Grubb, Brown (163). For Rad51 and Dmc1 co-staining immunofluorescence, polyclonal guinea pig α-Rad51 antibody serum and polyclonal goat α-Dmc1 antibody serum (a generous gift from D. Bishop, U. of Chicago) were diluted together 1:100 or 1:200 respectively into 1% BSA. 80 µl of diluted primary antibodies were added to each slide and covered with a 25 x 50 mm coverslip. Slides were incubated in a moist slide box at 4°C overnight (∼14-18 hrs). The next day, coverslips were removed in TBS and slides washed 2 X in TBS for 10 min. Donkey α- guinea pig Alexa 488 (Jackson ImmunoResearch 706-545-148) and donkey α-goat Alexa 594 (Fisher Cat#A11058) secondary antibodies were diluted 1:800 and 1:1000 respectively into 1% BSA. 80 µl of diluted secondary antibodies were added to each slide and covered with a coverslip. Slides were incubated in a moist slide box at 4°C for 2 hrs. Afterwards, slides were washed 2X in 1X TBS buffer for 10 min. Slides were air dried in the dark for 1 hr before 30 µL of DAPI was added to the slide. The slide was covered with a coverslip and sealed with nail polish. For nuclear spreads, microscopy images were taken using a Zeiss Axio Imager.Z2 microscope with a Zeiss Plan-Apochromat 100X objective.

### Fluorescent spore assays

Images of the fluorescent spores in tetrads used to score chromosome VIII nondisjunction and crossovers were taken using a Zeiss Axio Imager.Z2 microscope with a Zeiss Plan-Apochromat 63X objective. Cells were sporulated on solid Spo at 30°C for at least 3 days or in liquid Spo at 30°C for at least 24 hours before scoring. The genetic configurations of fluorescent markers for each system and patterns of fluorescent spore predicted are shown in Figs S2 and S4.

In the RGC system, parental ditypes (PD), tetratypes (T), and nonparental ditypes (NPD) can be distinguished between the *CEN8-ARG4* and *ARG4-THR1* intervals based on resulting fluorescent spore patterns (Fig S4). However, an NPD in the *CEN8-ARG4* interval results in the same fluorescent spore pattern as Chr VIII nondisjunction (Fig S4), making these two events indistinguishable. Therefore, the number of NPD tetrads in the *CEN8-ARG4* interval was estimated using the frequency of chromosome VIII nondisjunction observed in the RCEN system. This MI nondisjunction frequency was multiplied by the total number of RGC tetrads and that number was subtracted from the white:white;black;black tetrads to get the NPDs. For WT, no correction was done because no Chr VIII nondisjunction was observed in this strain. For *pif1-md*, *hedΔR* and *NDT80-mid*, the number of *CEN8-ARG4* NPD tetrads that could be explained by the frequency of chr VIII nondisjunction in the RCEN system was subtracted during map distance calculations. For *hedΔR NDT80-mid*, *hed1ΔR pif1-md* and *hed1Δ NDT80-mid pfi1-md*, all of *CEN8-ARG4* non-parental ditype tetrads were assumed to be due to nondisjunction [122]. Genetic map distance was calculated using the Perkins equation [164] using the Stahl lab online tool (https://elizabethhousworth.com/StahlLabOnlineTools/). Genetic interference ratios between *CEN8-ARG4* and *ARG4-THR1* intervals were calculated as in Malkova, Swanson (137) using the Stahl lab online tool. One interval was designated as the reference and tetrads from this interval were divided into two groups: one with parental ditypes (PDs) that have no crossovers and the other with tetratypes (Ts) or nonparental ditypes (NPDs) that contain at least one crossover. Map distances in the adjacent test interval were then calculated using tetrads from the PD or T+NPD groups from the reference interval. The map distance in the test interval derived from the T+NPD group was divided by that obtained from the PD group to give the crossover interference ratio. If crossovers in the reference interval interfered with crossovers in the test interval, this ratio should be less than one, while the absence of interference gives a ratio equal to one. Significance of crossover interference between the *CEN8-ARG4* and *ARG4-THR1* intervals was done using a G-test of independence.

### Physical Analysis of crossovers, noncrossovers and dHJs

To analyze crossover and noncrossover formation at the *HIS4-LEU2* DSB hotspot in meiotic timecourses [123]. Genomic DNA was isolated from sporulation cultures using MasterPure Yeast DNA Purification kit (Lucigen MPY80200). Processing of genomic DNA samples and Southern blot analysis used to visualize crossovers and noncrossovers at the *HIS4LEU2* hotspot were done as in [165]. For detection of joint molecules including SEIs, IH-dHJs and IS-dHJs, cells from meiotic time courses were subjected to psoralen-crosslinking, followed by extraction of genomic DNA, restriction digest with XhoI and 2D gel Southern blot analysis as previously described in Sandhu, Monge Neria (127).

### *URA3-tel-ARG4* hotspot analysis

To prevent mutations from accumulating during vegetative growth due to a lack of mismatch repair in the NH2700 *hedΔR msh2Δ* diploid, the haploid parents were first streaked out from freezer stocks onto YPDcom and allowed to grow for 51 hours at 30°C. Five single colonies from each haploid were combined in 200 μL YPD in a microfuge tube and the cells thoroughly mixed by vortexing. The entire volume was pipetted onto a YPD plate and the cells incubated at 30°C for 7 hours to allow mating. Cells were scraped from the plate, patched onto a plate containing sporulation medium (1% potassium acetate, 2% agar and 0.0004% histidine) and incubated for two days at 30°C. The resulting tetrads were dissected onto YPDcom.

Spore colonies from tetrads that exhibited four viable spores and 2:2 segregation for *hphMX* and *natMX4* were inoculated into 15 ml glass test tubes containing 2 ml YPD and incubated overnight at 30°C with shaking. Care was taken to ensure that the entire spore colony was used for each inoculation. For storage, 0.4 ml from each culture was transferred to a well in a sterile 1 ml 96 well plate containing 0.4 ml 50% glycerol and frozen at −80°C. In addition, 0.1 ml cell culture was transferred to the corresponding positions in a 0.45 ml 96 well plate with pointy bottoms (Fisher Scientific, #249946). Cells were pelleted in a Beckman tabletop centrifuge at 3000 rpm for 10 minutes and the supernatants were removed by aspiration. Each plate was sealed using a Microseal “F” PCR plate seal (Bio-Rad, #MSF1001) and stored at −80°C. This process was repeated to generate nine plates containing spore colonies from a total of 216 tetrads (9 * 24 tetrad/plate).

DNA was isolated from the 0.1 ml cell pellets as follows. Two 96 well plates were processed at a time. First, 5.5 ml Buffer 1 (100 ml 1 M sorbitol, 0.1 M EDTA) was placed in a 100 ml sterile reservoir (Stellar Scientific, #1930-1030) to which 1430 μL 10 mg/ml Zymolyase 100T (100 units/mg)(US Biological, #Z1004) was added. To spheroplast the cells, each cell pellet was resuspended in 20 μL of Buffer 1 + Zymolyase using a multichannel pipettor. The plates were covered and incubated with shaking for 30 min at 37°C. Six ml of Buffer 2 (50 mM Tris-HCl, pH 8, 20 mM EDTA, 0.35 M SDS) was placed in a fresh reservoir and 20 μl was added to each well and mixed well with a pipettor. The plates were covered and incubated for 30 min at 65°C with rotation. Six ml 5 M postassium acetate in a reservoir was used to distribute 16 μL to each well. After mixing well, the plates were covered and incubated at 4°C for 10 min. Nine ml 1 X TE (10 mM Tris-HCl, pH8, 1 mM EDTA) was used to add 80 μL TE to each well, after which the plates were covered and spun in a Beckman tabletop centrifuge for 60 min at 3500 rpm at 4°C. To precipitate the DNA, 50 μL supernatant from was transferred to a new 96 well plate containing 50 ul isopropanol in each well and mixed. The plates were covered with a plastic lid and incubated at room temperature for five minutes. DNA pellets were obtained by spinning the plates using a Beckman tabletop centrifuge for 10 min at 3500 rpm, after which the supernatants were carefully removed by aspiration. The DNA pellets were washed with 100 μl 70% ethanol and centrifuged as before. After removing the supernatants by aspiration, uncovered plates were placed at 42°C for 30 minutes to allow the pellets to dry. After the addition of 30 μL TE each well and the plates were sealed and incubated at 4°C overnight. The next day the DNA was resuspended by mixing with the pipettor. The DNA concentration of 1 μL of each sample was determined using the Qubit dsDNA HS Assay kit (Fisher Scientific, #33231) in black, round bottom non-treated 96 well plates (Fisher Scientific 07200762) per the manufacturer’s instructions using a BioTek Synergy 2 fluorimeter.

To facilitate the DNA sequencing analysis PCR amplicons containing the 6,865 bp recombination interval were amplified using the following primers: ACGGCACCACTATAAACCCG and GTGGGCTAAAGAACGCGAAC with spore specific barcodes on the 5’ end (S2_Data). All reactions were performed in 96 well plates as indicated in S2_Data. Each reaction contained 20 μL and used Q5 polymerase (New England Biolabs, #M0491). The reagents were distributed into each well of a 96 well plate as follows. A “Column Master Mix” was created by adding (13 X 4.0 μL 5X Q5 reaction buffer) + (13 X 2.6 μL dNTPs) + (13 X 0.75 μL water) + (13 X 0.4 μL 50 mM MgCl_2_) + (13 X 0.5 μL Q5 polymerase) and 82 μL was added to each of 12 strips containing eight 0.2 ml tubes. Ten μL of 10 uM Forward primer for the appropriate column was then aliquoted into each tube of a given strip. After mixing, 9.25 μL from each tube in a single strip was aliquoted into the appropriate column of a 96 well PCR plate. A “Row Master Mix” was generated by adding 14 X 8.75 μL water and then aliquoting 122.5 μL water into 8 strips of 8 tubes. Each strip represents a row, so 14 μL of the 10 μM Reverse primer designated for that row was added to each tube within a strip. 9.75 μL of the Row Master mix was then added to the 12 wells in each row of the PCR plate already containing the Column mix. Finally, one μL of DNA was added to the appropriate well of each plate. The plate was covered with an optically clear PCR plate plastic seal and spun for 2 min at 2000 rpm in a Beckman tabletop centrifuge. PCR reactions were performed using a Roche LightCycler 96 with the following program: 98°C, 40 sec; 98°C 20 sec, 67°C, 20 sec, 72°C, 7 min (30 cycles); 72°C, 10 min; 20°C hold.

Once the PCRs were complete, the samples were centrifuged as before to collect all of the liquid at the bottom of each well. Two μL from each reaction were run on a 0.9% agarose gel in 0.5X TBE buffer at 100 V for one hour. All reactions were examined for the presence of an ∼6 kb fragment. There were 159 tetrads in which all four spores exhibited a prominent PCR band and were included for the sequencing analysis. Four μL from each successful reaction were combined into one 1.5 ml microfuge tube and the primers were removed using Agencourt AMPure™ XP beads (Beckman Coulter #A63881).

### DNA sequencing and analysis

The pooled amplicons were sequenced on a Pacbio RS II instrument. Subsequently, barcodes were identified among PacBio reads (2.53 million) using the locate function of seqkit (Shen et al., 2016). Based on the barcode information the Pacbio-fastq file (https://www.ncbi.nlm.nih.gov/sra/PRJNA873686) was demultiplexed into sample specific fastq files (n=794) using a custom script. Each of the sample specific fastq files was aligned to the hotspot (Ahuja et al., 2021) using minimap2 [166], Base composition per nucleotide per read was determined using the sam2tsv function of the jvarkit [167].

Subsequently, phase was determined by a custom script that functions as follows: (1) determine the base(s) present at each polymorphic position (n=48) on each read (n= 2,384,252); (2) assign W (WT) or S (SNPs) for each polymorphic position per read depending on the nucleotide(s) present at the position; (3) select reads that have information on all polymorphic loci; and (4) group reads by identical marker configuration at all polymorphic locations and plot them as haplotype and number of reads that support the haplotype. Next, these plots were manually analyzed, and strands were placed into the following categories: (1) only one haplotype, signifying homoduplex at all markers across the interval i.e., the two strands are identical; (2) two haplotypes across the interval signifies two strands of the spore colony; (3) more than two haplotypes in these plots. In these cases, two haplotypes are supported by a majority of reads and some other reads may be promiscuous rearrangement due to PCR issues and these need to be manually inspected and removed. The final haplotypes used in the current project are derived from 1,179,721 reads, where per haplotype reads have a mean of 1584 reads with standard deviation of 1286 reads. Once, a consensus haplotype for each strand of each spore colony is determined and is scored in S2 Data. Schematics of the marker configurations in each tetrad are in S3 Data.

### Supporting information

**S1 Data.** Contains the data and calculations used for all of the numerical data presented in figures. (XLXS)

**S2 Data.** Contains the data used for the *URA3-tel-ARG4* hotspot analysis comparing WT and *hedΔR*. (XLXS)

**S3 Data.** Contains schematics of the distribution of polymorphisms in the four chromatids of each tetrad. (HTML)

**S1 Table.** Strains. (PDF)

**S2 Table.** Plasmids (PDF)

## Acknowledgements

We are grateful to Doug Bishop, Brooks Crickard, Bruce Futcher, Wolf Heyer, Scott Keeney, Ed Luk, Aaron Neiman, Adam Rosebrock and members of the Hollingsworth and Luk labs for helpful discussions. Bob Galione, Wolf Heyer and Aaron Neiman provided helpful comments on the manuscript. Scott Keeney and Allison Marullo provided plasmids and Eric Greene shared unpublished data with us. We thank Doug Bishop and Jennifer Grubb for antibodies and helpful technical advice and Aaron Neiman and Ed Luk for allowing us access to a fluorescence microscope and plate reader, respectively. We appreciate the help of Lihong Wan and Bob Gaglione with tetrad dissection and/or plasmid/strain construction. We are grateful to the Center for Cancer Research Sequencing Facility for high-throughput sequencing. This work used resources at the National Institutes of Health (NIH) Biowulf cluster (http://hpc.nih.gov).

## SUPPLEMENTAL FIGURES

**Fig S1. Validation of Dmc1, Rad51 and Red1 antibodies for cytological analyses.** (A) Detection of Dmc1 and Rad51 foci. Chromosome spreads at the indicated timepoints in Spo medium made from WT (NH716), *dmc1Δ* (NH2664), or *rad51Δ* (NH793) were probed with antibodies against either Rad51 or Dmc1 and detected by indirect immunofluorescence. DAPI staining was used to detect DNA. Scale bar is 2 µm. (B) Whole cell Red1 immunofluorescence. The diploids *ndt80Δ* (NH2234) and *ndt80Δ red1Δ* (NH2233) were incubated in Spo medium for the indicated times. Cells were fixed, stained with DAPI and probed with α-Red1 antibodies. Red1 was detected by indirect immunofluorescence.

**Fig S2. Fluorescent spore assays for measuring MI non-disjunction, map distance and interference (adapted from Thacker et al. 2011)** (A) RCEN. The *RFP* and *CFP* genes are localized at allelic positions adjacent to the Chromosome VIII centromere. This configuration was used to measure chromosome VIII nondisjunction. Examples of tetrads with normal segregation and MI nondisjunction are shown. (B) RGC. The *RFP, GFP\** and *CFP* genes are localized at different positions on Chromosome VIII. This configuration is used to measure map distances and interference between *CEN8*-*ARG4* and *ARG4-THR1*. Representative examples of fluorescent spore patterns for parental ditype and tetratype asci for both intervals are shown. Note that non-parental ditypes formed between *CEN8* and *ARG4* exhibit the same spore pattern as MI nondisjunction. A more complete depiction of the various fluorescent spore patterns is shown in Fig S4.

**Fig S3. Crossover and noncrossover formation at the *HIS4LEU2* hotspot in various mutant diploids.** (A) Southern blot of one of two timecourses used for this figure containing WT, *hedΔR*, *hedΔR NDT80-mid*, *pif1-md* (NH2657), *hedΔR pif1-md* (NH2691) and *hedΔR NDT80-mid pif1-md* (NH2661). Genomic DNA was digested with XhoI and NgoMIV to detect the CO2 and NCO1-specific bands described in Fig 4A. (B) Quantification of CO2 showing the average values from two different timecourses. (C) Quantification of NCO1 showing the average values from two different timecourses. Note that one of the *hedΔR NDT80-mid* replicates exhibited an unusually high value at the six hour timepoint which skewed the result.

**Fig S4. Possible fluorescent spore patterns for tetrads from an RGC diploid**. The left side shows the positions of crossovers (indicated by an “X”) between *CEN8::RFP* and *ARG4::GFP\** (Interval 1) and *ARG4::GFP\** and *THR1::CFP* (Interval 2). The next column shows the configuration of fluorescent protein genes on each chromatid after the indicated crossovers. The table indicates whether a given interval is a parental ditype (PD), tetratype (T) or nonparental ditype (NPD). The tetrad indicated in the gray shaded part of the diagram can be due either to an NPD in Interval 1 or MI nondisjunction. R = red, A = aqua, W = white, Bl = black, P = purple, G = green, Y = yellow, B = blue

**Figure S5. Gene conversion tracts in *hedΔR* and WT** Gene conversion tracts in noncrossovers (A) and crossovers (B) in *hedΔR* (top) and wild type (bottom). Thick colored bars and thin gray bars indicate minimum and maximum NMS tracts, respectively. Turquoise—heteroduplex on one chromatid; brown—heteroduplex on two chromatids. Vertical axis–tetrad identifiers; vertical lines— marker positions. Underlying data are in Data_S2, sheet 6 and in [126].

**Figure S6. Complementation of different alleles of *RRM3* and *PIF1* in the *hed****Δ**R NDT80-mid*** **background.** (A) Spore viability. Single colonies of the *hedΔR NDT80-mid rrm3Δ* diploid (NH2596) containing two copies of the vector (v) (pRS306), *RRM3* (pBG22) or *P_REC8_-RRM3* (pJW7) and the *hedΔR NDT80-mid pif1-md* diploid, NH2661, containing two copies of pRS306, *PIF1* (pJW5), *pif1-m1* (pJW5-m1), *PIF1-3FLAG* (pJW11) or *pif1-m1-3FLAG* (pJW14) were sporulated on solid medium and at least 20 tetrads were dissected per transformant. The statistical significance of differences between strains was determined using the Mann-Whitney test (* = *p*<.05; ** = *p*<.01; *** = *p*<.001, **** = *p*<.0001). (B) Distribution of viable spores in tetrads for a subset of the *hedΔR NDT80-mid pif1-md* dissections shown in Panel A. (C) Distribution of viable spores in tetrads for the *hedΔR NDT80-mid rrm3Δ* dissections shown in Panel A.

**Figure S7. *PIF1* promotes meiotic progression when Rad51 is constitutively active during meiotic prophase I.** (A) Meiotic progression. Timecourse analysis to determine the percentage of bi and tetranucleate cells was performed as in Figure 1A and the average %MI + MII values plotted for each timepoint. The strains and data for WT, *hedΔR*, and *hedΔR NDT80-mid* are the same as in Figure 3A with the addition of two biological replicates. The new strains were *pif1-md* (NH2657) (*n* = 7), *hedΔR pif1-md* (NH2691) (*n* = 11) and *hedΔR NDT80-mid pif1-md* (NH2661) (*n* = 7). (B) T_50_ values calculated for the timecourses in Panel A. (C) Prophase lifespan. The data for WT, *hedΔR* and *hedΔR NDT80-mid* for Panels B and C were taken from Figure 3. Data for *pif1-md* containing strains is from the timecourses in Panel A.

## Notes

### Competing Interest Statement

The authors have declared no competing interest.

## REFERENCES

1. Petronczki M, Siomos MF, Nasmyth K. *Un menage a quatre*: the molecular biology of chromosome segregation in meiosis. Cell. 2003;112(4):423–40.

2. Hassold T, Hunt P. To err (meiotically) is human: the genesis of human aneuploidy. Nat Rev Genet. 2001;2(4):280–91.

3. Villeneuve AM, Hillers KJ. Whence meiosis? Cell. 2001;106(6):647–50.

4. Keeney S, Giroux CN, Kleckner N. Meiosis-specific DNA double-strand breaks are catalyzed by Spo11, a member of a widely conserved protein family. Cell. 1997;88(3):375–84.

5. Borde V, Goldman AS, Lichten M. Direct coupling between meiotic DNA replication and recombination initiation. Science. 2000;290(5492):806–9.

6. Lam I, Keeney S. Mechanism and regulation of meiotic recombination initiation. Cold Spring Harb Perspect Biol. 2014;7(1):a016634.

7. Pan J, Sasaki M, Kniewel R, Murakami H, Blitzblau HG, Tischfield SE, et al. A hierarchical combination of factors shapes the genome-wide topography of yeast meiotic recombination initiation. Cell. 2011;144(5):719–31.

8. Garcia V, Phelps SE, Gray S, Neale MJ. Bidirectional resection of DNA double-strand breaks by Mre11 and Exo1. Nature. 2011;479(7372):241–4.

9. Sun H, Treco D, Szostak JW. Extensive 3’-overhanging, single-stranded DNA associated with the meiosis-specific double-strand breaks at the *ARG4* recombination initiation site. Cell. 1991;64(6):1155–61.

10. Mimitou EP, Yamada S, Keeney S. A global view of meiotic double-strand break end resection. Science. 2017;355(6320):40–5.

11. Bishop DK, Park D, Xu L, Kleckner N. *DMC1*: a meiosis-specific yeast homolog of *E. coli recA* required for recombination, synaptonemal complex formation, and cell cycle progression. Cell. 1992;69(3):439–56.

12. Shinohara A, Ogawa H, Ogawa T. Rad51 protein involved in repair and recombination in *S. cerevisiae* is a RecA-like protein. Cell. 1992;69(3):457–70.

13. Habu T, Taki T, West A, Nishimune Y, Morita T. The mouse and human homologs of *DMC1*, the yeast meiosis-specific homologous recombination gene, have a common unique form of exon-skipped transcript in meiosis. Nucleic Acids Res. 1996;24(3):470–7.

14. Yoshida K, Kondoh G, Matsuda Y, Habu T, Nishimune Y, Morita T. The mouse RecA-like gene Dmc1 is required for homologous chromosome synapsis during meiosis. Mol Cell. 1998;1(5):707–18.

15. Sung P. Catalysis of ATP-dependent homologous DNA pairing and strand exchange by yeast *RAD51* protein. Science. 1994;265(5176):1241–3.

16. Hong EL, Shinohara A, Bishop DK. *Saccharomyces cerevisiae* Dmc1 protein promotes renaturation of single-strand DNA (ssDNA) and assimilation of ssDNA into homologous super-coiled duplex DNA. J Biol Chem. 2001;276(45):41906–12.

17. Brown MS, Bishop DK. DNA strand exchange and RecA homologs in meiosis. Cold Spring Harb Perspect Biol. 2014;7(1):a016659.

18. Sheridan S, Bishop DK. Red-Hed regulation: recombinase Rad51, though capable of playing the leading role, may be relegated to supporting Dmc1 in budding yeast meiosis. Genes Dev. 2006;20(13):1685–91.

19. Brown MS, Grubb J, Zhang A, Rust MJ, Bishop DK. Small Rad51 and Dmc1 complexes often co-occupy both ends of a meiotic DNA double strand break. PLoS Genet. 2015;11(12):e1005653.

20. Lee MH, Chang YC, Hong EL, Grubb J, Chang CS, Bishop DK, et al. Calcium ion promotes yeast Dmc1 activity via formation of long and fine helical filaments with single-stranded DNA. J Biol Chem. 2005;280(49):40980–4.

21. Ogawa T, Yu X, Shinohara A, Egelman EH. Similarity of the yeast *RAD51* filament to the bacterial RecA filament. Science. 1993;259(5103):1896–9.

22. Hinch AG, Becker PW, Li T, Moralli D, Zhang G, Bycroft C, et al. The configuration of RPA, *RAD51*, and *DMC1* binding in meiosis reveals the nature of critical recombination intermediates. Mol Cell. 2020;79(4):689–701 e10.

23. Benson FE, Stasiak A, West SC. Purification and characterization of the human Rad51 protein, an analogue of *E. coli* RecA. EMBO J. 1994;13(23):5764–71.

24. Sheridan SD, Yu X, Roth R, Heuser JE, Sehorn MG, Sung P, et al. A comparative analysis of Dmc1 and Rad51 nucleoprotein filaments. Nucleic Acids Res. 2008;36(12):4057–66.

25. Cloud V, Chan YL, Grubb J, Budke B, Bishop DK. Rad51 is an accessory factor for Dmc1-mediated joint molecule formation during meiosis. Science. 2012;337(6099):1222–5.

26. Lao JP, Oh SD, Shinohara M, Shinohara A, Hunter N. Rad52 promotes postinvasion steps of meiotic double-strand-break repair. Mol Cell. 2008;29(4):517–24.

27. Bishop DK. RecA homologs Dmc1 and Rad51 interact to form multiple nuclear complexes prior to meiotic chromosome synapsis. Cell. 1994;79(6):1081–92.

28. Kurzbauer MT, Uanschou C, Chen D, Schlogelhofer P. The recombinases *DMC1* and *RAD51* are functionally and spatially separated during meiosis in *Arabidopsis*. Plant Cell. 2012;24(5):2058–70.

29. Lan WH, Lin SY, Kao CY, Chang WH, Yeh HY, Chang HY, et al. Rad51 facilitates filament assembly of meiosis-specific Dmc1 recombinase. Proc Natl Acad Sci U S A. 2020;117(21):11257–64.

30. Da Ines O, Degroote F, Goubely C, Amiard S, Gallego ME, White CI. Meiotic recombination in *Arabidopsis* is catalysed by DMC1, with RAD51 playing a supporting role. PLoS Genet. 2013;9(9):e1003787.

31. Petukhova G, Stratton S, Sung P. Catalysis of homologous DNA pairing by yeast Rad51 and Rad54 proteins. Nature. 1998;393(6680):91–4.

32. Nimonkar AV, Dombrowski CC, Siino JS, Stasiak AZ, Stasiak A, Kowalczykowski SC. *Saccharomyces cerevisiae* Dmc1 and Rad51 proteins preferentially function with Tid1 and Rad54 proteins, respectively, to promote DNA strand invasion during genetic recombination. J Biol Chem. 2012;287(34):28727–37.

33. Bianchi M, DasGupta C, Radding CM. Synapsis and the formation of paranemic joints by *E. coli* RecA protein. Cell. 1983;34(3):931–9.

34. Riddles PW, Lehman IR. The formation of paranemic and plectonemic joints between DNA molecules by the *recA* and single-stranded DNA-binding proteins of *Escherichia coli*. J Biol Chem. 1985;260(1):165–9.

35. Van Komen S, Petukhova G, Sigurdsson S, Stratton S, Sung P. Superhelicity-driven homologous DNA pairing by yeast recombination factors Rad51 and Rad54. Mol Cell. 2000;6(3):563–72.

36. Christiansen G, Griffith J. Visualization of the paranemic joining of homologous DNA molecules catalyzed by the RecA protein of *Escherichia coli*. Proc Natl Acad Sci U S A. 1986;83(7):2066–70.

37. Wright WD, Heyer WD. Rad54 functions as a heteroduplex DNA pump modulated by its DNA substrates and Rad51 during D loop formation. Mol Cell. 2014;53(3):420–32.

38. Shibata T, DasGupta C, Cunningham RP, Radding CM. Purified *Escherichia coli recA* protein catalyzes homologous pairing of superhelical DNA and single-stranded fragments. Proc Natl Acad Sci U S A. 1979;76(4):1638–42.

39. Petukhova G, Van Komen S, Vergano S, Klein H, Sung P. Yeast Rad54 promotes Rad51-dependent homologous DNA pairing via ATP hydrolysis-driven change in DNA double helix conformation. J Biol Chem. 1999;274(41):29453–62.

40. Piazza A, Shah SS, Wright WD, Gore SK, Koszul R, Heyer WD. Dynamic processing of displacement loops during recombinational DNA repair. Mol Cell. 2019;73(6):1255–66 e4.

41. Heyer WD, Li X, Rolfsmeier M, Zhang XP. *Rad54*: the Swiss Army knife of homologous recombination? Nucleic Acids Res. 2006;34(15):4115–25.

42. Crickard JB. Discrete roles for Rad54 and Rdh54 during homologous recombination. Curr Opin Genet Dev. 2021;71:48–54.

43. Crickard JB, Moevus CJ, Kwon Y, Sung P, Greene EC. Rad54 drives ATP hydrolysis-dependent DNA sequence alignment during homologous recombination. Cell. 2020;181(6):1380–94 e18.

44. Crickard JB, Kwon Y, Sung P, Greene EC. Rad54 and Rdh54 occupy spatially and functionally distinct sites within the Rad51-ssDNA presynaptic complex. EMBO J. 2020;39(20):e105705.

45. Tavares EM, Wright WD, Heyer WD, Le Cam E, Dupaigne P. *In vitro* role of Rad54 in Rad51-ssDNA filament-dependent homology search and synaptic complexes formation. Nat Commun. 2019;10(1):4058.

46. Mazin AV, Bornarth CJ, Solinger JA, Heyer WD, Kowalczykowski SC. Rad54 protein is targeted to pairing loci by the Rad51 nucleoprotein filament. Mol Cell. 2000;6(3):583–92.

47. Clever B, Interthal H, Schmuckli-Maurer J, King J, Sigrist M, Heyer WD. Recombinational repair in yeast: functional interactions between Rad51 and Rad54 proteins. EMBO J. 1997;16(9):2535–44.

48. Jiang H, Xie Y, Houston P, Stemke-Hale K, Mortensen UH, Rothstein R, et al. Direct association between the yeast Rad51 and Rad54 recombination proteins. J Biol Chem. 1996;271(52):33181–6.

49. Mazin AV, Alexeev AA, Kowalczykowski SC. A novel function of Rad54 protein: stabilization of the Rad51 nucleoprotein filament. J Biol Chem. 2003;278(16):14029–36.

50. Li X, Heyer WD. *RAD54* controls access to the invading 3’-OH end after *RAD51*-mediated DNA strand invasion in homologous recombination in *Saccharomyces cerevisiae*. Nucleic Acids Res. 2009;37(2):638–46.

51. Solinger JA, Kiianitsa K, Heyer WD. Rad54, a Swi2/Snf2-like recombinational repair protein, disassembles Rad51:dsDNA filaments. Mol Cell. 2002;10(5):1175–88.

52. Maloisel L, Fabre F, Gangloff S. DNA polymerase delta is preferentially recruited during homologous recombination to promote heteroduplex DNA extension. Mol Cell Biol. 2008;28(4):1373–82.

53. Maloisel L, Bhargava J, Roeder GS. A role for DNA polymerase delta in gene conversion and crossing over during meiosis in *Saccharomyces cerevisiae*. Genetics. 2004;167(3):1133–42.

54. Li X, Stith CM, Burgers PM, Heyer WD. PCNA is required for initiation of recombination-associated DNA synthesis by DNA polymerase delta. Mol Cell. 2009;36(4):704–13.

55. Sneeden JL, Grossi SM, Tappin I, Hurwitz J, Heyer WD. Reconstitution of recombination-associated DNA synthesis with human proteins. Nucleic Acids Res. 2013;41(9):4913–25.

56. Allers T, Lichten M. Differential timing and control of noncrossover and crossover recombination during meiosis. Cell. 2001;106(1):47–57.

57. De Muyt A, Jessop L, Kolar E, Sourirajan A, Chen J, Dayani Y, et al. BLM helicase ortholog Sgs1 is a central regulator of meiotic recombination intermediate metabolism. Mol Cell. 2012;46(1):43–53.

58. Zakharyevich K, Tang S, Ma Y, Hunter N. Delineation of joint molecule resolution pathways in meiosis identifies a crossover-specific resolvase. Cell. 2012;149(2):334–47.

59. McMahill MS, Sham CW, Bishop DK. Synthesis-dependent strand annealing in meiosis. PLoS Biol. 2007;5(11):e299.

60. Kaur H, De Muyt A, Lichten M. Top3-Rmi1 DNA single-strand decatenase is integral to the formation and resolution of meiotic recombination intermediates. Mol Cell. 2015;57(4):583–94.

61. Tang S, Wu MKY, Zhang R, Hunter N. Pervasive and essential roles of the Top3-Rmi1 decatenase orchestrate recombination and facilitate chromosome segregation in meiosis. Mol Cell. 2015;57(4):607–21.

62. Bell L, Byers B. Occurrence of crossed strand-exchange forms in yeast DNA during meiosis. Proc Natl Acad Sci U S A. 1979;76(7):3445–9.

63. Schwacha A, Kleckner N. Identification of double Holliday junctions as intermediates in meiotic recombination. Cell. 1995;83(5):783–91.

64. Szostak JW, Orr-Weaver TL, Rothstein RJ, Stahl FW. The double-strand-break repair model for recombination. Cell. 1983;33(1):25–35.

65. Pyatnitskaya A, Borde V, De Muyt A. Crossing and zipping: molecular duties of the ZMM proteins in meiosis. Chromosoma. 2019;128(3):181–98.

66. Börner GV, Kleckner N, Hunter N. Crossover/noncrossover differentiation, synaptonemal complex formation, and regulatory surveillance at the leptotene/zygotene transition of meiosis. Cell. 2004;117(1):29–45.

67. Sym M, Engebrecht JA, Roeder GS. *ZIP1* is a synaptonemal complex protein required for meiotic chromosome synapsis. Cell. 1993;72(3):365–78.

68. Zickler D, Kleckner N. Recombination, pairing, and synapsis of homologs during meiosis. Cold Spring Harb Perspect Biol. 2015;7(6).

69. Grey C, de Massy B. Chromosome organization in early meiotic prophase. Front Cell Dev Biol. 2021;9:688878. doi: 10.3389/fcell.2021.688878.

70. Smith AV, Roeder GS. The yeast Red1 protein localizes to the cores of meiotic chromosomes. J Cell Biol. 1997;136(5):957–67.

71. Klein F, Mahr P, Galova M, Buonomo SBC, Michaelis C, Nairz K, et al. A central role for cohesions in sister chromatid cohesion, formation of axial elements, and recombination during yeast meiosis. Cell. 1999;98(1):91–103.

72. Hollingsworth NM, Goetsch L, Byers B. The *HOP1* gene encodes a meiosis-specific component of yeast chromosomes. Cell. 1990;61(1):73–84.

73. Humphryes N, Leung WK, Argunhan B, Terentyev Y, Dvorackova M, Tsubouchi H. The Ecm11-Gmc2 complex promotes synaptonemal complex formation through assembly of transverse filaments in budding yeast. PLoS Genet. 2013;9(1):e1003194.

74. Rockmill B, Roeder GS. A meiosis-specific protein kinase homolog required for chromosome synapsis and recombination. Genes Dev. 1991;5(12B):2392–404.

75. Hollingsworth NM, Gaglione R. The meiotic-specific Mek1 kinase in budding yeast regulates interhomolog recombination and coordinates meiotic progression with double-strand break repair. Curr Genet. 2019;65(3):631–41.

76. Leem SH, Ogawa H. The *MRE4* gene encodes a novel protein kinase homologue required for meiotic recombination in *Saccharomyces cerevisiae*. Nucleic Acids Res. 1992;20(3):449–57.

77. Xu L, Weiner BM, Kleckner N. Meiotic cells monitor the status of the interhomolog recombination complex. Genes Dev. 1997;11(1):106–18.

78. Niu H, Wan L, Baumgartner B, Schaefer D, Loidl J, Hollingsworth NM. Partner choice during meiosis is regulated by Hop1-promoted dimerization of Mek1. Mol Biol Cell. 2005;16(12):5804–18.

79. Kim KP, Weiner BM, Zhang L, Jordan A, Dekker J, Kleckner N. Sister cohesion and structural axis components mediate homolog bias of meiotic recombination. Cell. 2010;143(6):924–37.

80. Chen X, Suhandynata RT, Sandhu R, Rockmill B, Mohibullah N, Niu H, et al. Phosphorylation of the synaptonemal complex protein Zip1 regulates the crossover/noncrossover decision during yeast meiosis. PLoS Biol. 2015;13(12):e1002329.

81. Kadyk LC, Hartwell LH. Sister chromatids are preferred over homologs as substrates for recombinational repair in *Saccharomyces cerevisiae*. Genetics. 1992;132(2):387–402.

82. Bzymek M, Thayer NH, Oh SD, Kleckner N, Hunter N. Double Holliday junctions are intermediates of DNA break repair. Nature. 2010;464(7290):937–41.

83. Schwacha A, Kleckner N. Interhomolog bias during meiotic recombination: meiotic functions promote a highly differentiated interhomolog-only pathway. Cell. 1997;90(6):1123–35.

84. Callender TL, Laureau R, Wan L, Chen X, Sandhu R, Laljee S, et al. Mek1 down regulates Rad51 activity during yeast meiosis by phosphorylation of Hed1. PLoS Genet. 2016;12(8):e1006226.

85. Busygina V, Sehorn MG, Shi IY, Tsubouchi H, Roeder GS, Sung P. Hed1 regulates Rad51-mediated recombination via a novel mechanism. Genes Dev. 2008;22(6):786–95.

86. Tsubouchi H, Roeder GS. Budding yeast Hed1 down-regulates the mitotic recombination machinery when meiotic recombination is impaired. Genes Dev. 2006;20(13):1766–75.

87. Niu H, Wan L, Busygina V, Kwon Y, Allen JA, Li X, et al. Regulation of meiotic recombination via Mek1-mediated Rad54 phosphorylation. Mol Cell. 2009;36(3):393–404.

88. Lao JP, Cloud V, Huang CC, Grubb J, Thacker D, Lee CY, et al. Meiotic crossover control by concerted action of Rad51-Dmc1 in homolog template bias and robust homeostatic regulation. PLoS Genet. 2013;9(12):e1003978.

89. Hong S, Sung Y, Yu M, Lee M, Kleckner N, Kim KP. The logic and mechanism of homologous recombination partner choice. Mol Cell. 2013;51(4):440–53.

90. Bishop DK, Nikolski Y, Oshiro J, Chon J, Shinohara M, Chen X. High copy number suppression of the meiotic arrest caused by a *dmc1* mutation: *REC114* imposes an early recombination block and *RAD54* promotes a *DMC1*-independent DSB repair pathway. Genes Cells. 1999;4(8):425–44.

91. Chen X, Gaglione R, Leong T, Bednor L, de Los Santos T, Luk E, et al. Mek1 coordinates meiotic progression with DNA break repair by directly phosphorylating and inhibiting the yeast pachytene exit regulator Ndt80. PLoS Genet. 2018;14(11):e1007832.

92. Wu HY, Ho HC, Burgess SM. Mek1 kinase governs outcomes of meiotic recombination and the checkpoint response. Curr Biol. 2010;20(19):1707–16.

93. Hollingsworth NM. A new role for the synaptonemal complex in the regulation of meiotic recombination. Genes Dev. 2020;34(23-24):1562–4.

94. Prugar E, Burnett C, Chen X, Hollingsworth NM. Coordination of double strand break repair and meiotic progression in yeast by a Mek1-Ndt80 negative feedback loop. Genetics. 2017;206(1):497–512.

95. Subramanian VV, MacQueen AJ, Vader G, Shinohara M, Sanchez A, Borde V, et al. Chromosome synapsis alleviates Mek1-dependent suppression of meiotic DNA repair. PLoS Biol. 2016;14(2):e1002369.

96. Mu X, Murakami H, Mohibullah N, Keeney S. Chromosome-autonomous feedback down-regulates meiotic DNA break competence upon synaptonemal complex formation. Genes Dev. 2020;34(23-24):1605–18.

97. Lee MS, Higashide MT, Choi H, Li K, Hong S, Lee K, et al. The synaptonemal complex central region modulates crossover pathways and feedback control of meiotic double-strand break formation. Nucleic Acids Res. 2021;49(13):7537–53.

98. Okaz E, Arguello-Miranda O, Bogdanova A, Vinod PK, Lipp JJ, Markova Z, et al. Meiotic prophase requires proteolysis of M phase regulators mediated by the meiosis-specific APC/C Ama1. Cell. 2012;151(3):603–18.

99. Chu S, Herskowitz I. Gametogenesis in yeast is regulated by a transcriptional cascade dependent on Ndt80. Mol Cell. 1998;1(5):685–96.

100. Chu S, DeRisi J, Eisen M, Mulholland J, Botstein D, Brown PO, et al. The transcriptional program of sporulation in budding yeast. Science. 1998;282(5389):699–705.

101. Sourirajan A, Lichten M. Polo-like kinase Cdc5 drives exit from pachytene during budding yeast meiosis. Genes Dev. 2008;22(19):2627–32.

102. Clyne RK, Katis VL, Jessop L, Benjamin KR, Herskowitz I, Lichten M, et al. Polo-like kinase Cdc5 promotes chiasmata formation and cosegregation of sister centromeres at meiosis I. Nat Cell Biol. 2003;5(5):480–5.

103. Argunhan B, Leung WK, Afshar N, Terentyev Y, Subramanian VV, Murayama Y, et al. Fundamental cell cycle kinases collaborate to ensure timely destruction of the synaptonemal complex during meiosis. EMBO J. 2017;36(17):2488–509.

104. Hayashi M, Chin GM, Villeneuve AM. *C. elegans* germ cells switch between distinct modes of double-strand break repair during meiotic prophase progression. PLoS Genet. 2007;3(11):e191.

105. Toraason E, Horacek A, Clark C, Glover ML, Adler VL, Premkumar T, et al. Meiotic DNA break repair can utilize homolog-independent chromatid templates in *C. elegans*. Curr Biol. 2021;31(7):1508–14 e5.

106. Enguita-Marruedo A, Martin-Ruiz M, Garcia E, Gil-Fernandez A, Parra MT, Viera A, et al. Transition from a meiotic to a somatic-like DNA damage response during the pachytene stage in mouse meiosis. PLoS Genet. 2019;15(1):e1007439.

107. Liu Y, Gaines WA, Callender T, Busygina V, Oke A, Sung P, et al. Down-regulation of Rad51 activity during meiosis in yeast prevents competition with Dmc1 for repair of double-strand breaks. PLoS Genet. 2014;10(1):e1004005.

108. Davis AP, Symington LS. *RAD51*-dependent break-induced replication in yeast. Mol Cell Biol. 2004;24(6):2344–51.

109. Wilson MA, Kwon Y, Xu Y, Chung WH, Chi P, Niu H, et al. Pif1 helicase and Poldelta promote recombination-coupled DNA synthesis via bubble migration. Nature. 2013;502(7471):393–6.

110. Saini N, Ramakrishnan S, Elango R, Ayyar S, Zhang Y, Deem A, et al. Migrating bubble during break-induced replication drives conservative DNA synthesis. Nature. 2013;502(7471):389–92.

111. Buzovetsky O, Kwon Y, Pham NT, Kim C, Ira G, Sung P, et al. Role of the Pif1-PCNA vomplex in pol delta-dependent strand displacement DNA synthesis and break-induced replication. Cell Rep. 2017;21(7):1707–14.

112. Vernekar DV, Reginato G, Adam C, Ranjha L, Dingli F, Marsolier MC, et al. The Pif1 helicase is actively inhibited during meiotic recombination which restrains gene conversion tract length. Nucleic Acids Res. 2021;49(8):4522–33.

113. Cha RS, Weiner BM, Keeney S, Dekker J, Kleckner N. Progression of meiotic DNA replication is modulated by interchromosomal interaction proteins, negatively by Spo11p and positively by Rec8p. Genes Dev. 2000;14(4):493–503.

114. Lydall D, Nikolsky Y, Bishop DK, Weinert T. A meiotic recombination checkpoint controlled by mitotic checkpoint genes. Nature. 1996;383(6603):840–3.

115. Hochwagen A, Amon A. Checking your breaks: surveillance mechanisms of meiotic recombination. Curr Biol. 2006;16(6):R217–28.

116. Reitz D, Grubb J, Bishop DK. A mutant form of Dmc1 that bypasses the requirement for accessory protein Mei5-Sae3 reveals independent activities of Mei5-Sae3 and Rad51 in Dmc1 filament stability. PLoS Genet. 2019;15(12):e1008217.

117. Busygina V, Saro D, Williams G, Leung WK, Say AF, Sehorn MG, et al. Novel attributes of Hed1 affect dynamics and activity of the Rad51 presynaptic filament during meiotic recombination. J Biol Chem. 2012;287(2):1566–75.

118. Crickard JB, Kaniecki K, Kwon Y, Sung P, Lisby M, Greene EC. Regulation of Hed1 and Rad54 binding during maturation of the meiosis-specific presynaptic complex. EMBO J. 2018;37(7).

119. Cartagena-Lirola H, Guerini I, Manfrini N, Lucchini G, Longhese MP. Role of the *Saccharomyces cerevisiae* Rad53 checkpoint kinase in signaling double-strand breaks during the meiotic cell cycle. Mol Cell Biol. 2008;28(14):4480–93.

120. Cheslock PS, Kemp BJ, Boumil RM, Dawson DS. The roles of *MAD1*, *MAD2* and *MAD3* in meiotic progression and the segregation of nonexchange chromosomes. Nat Genet. 2005;37(7):756–60.

121. Hollingsworth NM, Ponte L, Halsey C. *MSH5*, a novel MutS homolog, facilitates meiotic reciprocal recombination between homologs in *Saccharomyces cerevisiae* but not mismatch repair. Genes Dev. 1995;9(14):1728–39.

122. Thacker D, Lam I, Knop M, Keeney S. Exploiting spore-autonomous fluorescent protein expression to quantify meiotic chromosome behaviors in *Saccharomyces cerevisiae*. Genetics. 2011;189(2):423–39.

123. Martini E, Diaz RL, Hunter N, Keeney S. Crossover homeostasis in yeast meiosis. Cell. 2006;126(2):285–95.

124. De Muyt A, Jessop L, Kolar E, Sourirajan A, Chen J, Dayani Y, et al. BLM helicase ortholog Sgs1 is a central regulator of meiotic recombination intermediate metabolism. Mol Cell. 2012;46:43–53.

125. Marsolier-Kergoat M-C, Khan MM, Schott J, Zhu X, Llorente B. Mechanistic view and genetic control of DNA recombination during meiosis. Mol Cell. 2018;70:9–20.

126. Ahuja JS, Harvey CS, Wheeler DL, Lichten M. Repeated strand invasion and extensive branch migration are hallmarks of meiotic recombination. Mol Cell. 2021;81(20):4258–70 e4.

127. Sandhu R, Monge Neria F, Monge Neria J, Chen X, Hollingsworth NM, Börner GV. DNA helicase Mph1(FANCM) ensures meiotic recombination between parental chromosomes by dissociating precocious displacement loops. Dev Cell. 2020;53(4):458–72 e5.

128. Cooper TJ, Crawford MR, Hunt LJ, Marsolier-Kergoat M-C, Llorente B, Neale MJ. Mismatch repair impedes meiotic crossover interference. bioRxiv. 2018. doi: 10.1101/480418.

129. Martini E, Borde V, Legendre M, Audic S, Regnault B, Soubigou G, et al. Genome-wide analysis of heteroduplex DNA in mismatch repair-deficient yeast cells reveals novel properties of meiotic recombination pathways. PLoS Genet. 2011;7:e1002305.

130. Oke A, Anderson CM, Yam P, Fung JC. Controlling meiotic recombinational repair - specifying the roles of ZMMs, Sgs1 and Mus81/Mms4 in crossover formation. PLoS Genet. 2014;10:e1004690.

131. Lee BH, Amon A. Role of Polo-like kinase *CDC5* in programming meiosis I chromosome segregation. Science. 2003;300(5618):482–6.

132. Grandin N, Reed SI. Differential function and expression of *Saccharomyces cerevisiae* B-type cyclins in mitosis and meiosis. Mol Cell Biol. 1993;13(4):2113–25.

133. Ivessa AS, Zhou JQ, Zakian VA. The *Saccharomyces* Pif1p DNA helicase and the highly related Rrm3p have opposite effects on replication fork progression in ribosomal DNA. Cell. 2000;100(4):479–89.

134. Lahaye A, Stahl H, Thines-Sempoux D, Foury F. *PIF1*: a DNA helicase in yeast mitochondria. EMBO J. 1991;10(4):997–1007.

135. Schulz VP, Zakian VA. The *Saccharomyces PIF1* DNA helicase inhibits telomere elongation and *de novo* telomere formation. Cell. 1994;76(1):145–55.

136. Sturtevant AH. The linear arrangement of six X-linked factors in *Drosophila*, as shown by their mode of association. Journal of Experimental Zoology. 1913;(14):43–59.

137. Malkova A, Swanson J, German M, McCusker JH, Housworth EA, Stahl FW, et al. Gene conversion and crossing over along the 405-kb left arm of *Saccharomyces cerevisiae* chromosome VII. Genetics. 2004;168(1):49–63.

138. Azvolinsky A, Dunaway S, Torres JZ, Bessler JB, Zakian VA. The *S. cerevisiae* Rrm3p DNA helicase moves with the replication fork and affects replication of all yeast chromosomes. Genes Dev. 2006;20(22):3104–16.

139. Bochman ML, Sabouri N, Zakian VA. Unwinding the functions of the Pif1 family helicases. DNA Repair (Amst). 2010;9(3):237–49.

140. Claussin C, Vazquez J, Whitehouse I. Single-molecule mapping of replisome progression. Mol Cell. 2022;82(7):1372–82 e4.

141. Muellner J, Schmidt KH. Yeast Genome Maintenance by the Multifunctional *PIF1* DNA Helicase Family. Genes (Basel). 2020;11(2).

142. Chen CF, Pohl TJ, Pott S, Zakian VA. Two Pif1 family DNA helicases cooperate in centromere replication and segregation in *Saccharomyces cerevisiae*. Genetics. 2019;211(1):105–19.

143. Tran PLT, Pohl TJ, Chen CF, Chan A, Pott S, Zakian VA. *PIF*1 family DNA helicases suppress R-loop mediated genome instability at tRNA genes. Nat Commun. 2017;8:15025.

144. Ho HC, Burgess SM. Pch2 acts through Xrs2 and Tel1/ATM to modulate interhomolog bias and checkpoint function during meiosis. PLoS Genet. 2011;7(11):e1002351.

145. Carballo JA, Johnson AL, Sedgwick SG, Cha RS. Phosphorylation of the axial element protein Hop1 by Mec1/Tel1 ensures meiotic interhomolog recombination. Cell. 2008;132(5):758–70.

146. MacQueen AJ, Hochwagen A. Checkpoint mechanisms: the puppet masters of meiotic prophase. Trends Cell Biol. 2011;21(7):393–400.

147. Crawford MR, Cooper TJ, Marsolier-Kergoat M, Llorente B, Neale MJ. Separable roles of the DNA damage response kinase Mec1^ATR^ and its activator Rad2^RAD17^ within the regulation of meiotic recombination. bioRxiv. 2018. doi: 10.1101/496182.

148. Goldfarb T, Lichten M. Frequent and efficient use of the sister chromatid for DNA double-strand break repair during budding yeast meiosis. PLoS Biol. 2010;8(10):e1000520.

149. Chen YK, Leng CH, Olivares H, Lee MH, Chang YC, Kung WM, et al. Heterodimeric complexes of Hop2 and Mnd1 function with Dmc1 to promote meiotic homolog juxtaposition and strand assimilation. Proc Natl Acad Sci U S A. 2004;101(29):10572–7.

150. Zickler D, Espagne E. *Sordaria*, a model system to uncover links between meiotic pairing and recombination. Semin Cell Dev Biol. 2016;54:149–57.

151. Arbel A, Zenvirth D, Simchen G. Sister chromatid-based DNA repair is mediated by *RAD54*, not by *DMC1* or *TID1*. EMBO J. 1999;18(9):2648–58.

152. Sanchez H, Kertokalio A, van Rossum-Fikkert S, Kanaar R, Wyman C. Combined optical and topographic imaging reveals different arrangements of human *RAD54* with presynaptic and postsynaptic *RAD51*-DNA filaments. Proc Natl Acad Sci U S A. 2013;110(28):11385–90.

153. Dai J, Voloshin O, Potapova S, Camerini-Otero RD. Meiotic knockdown and complementation reveals essential role of *RAD51* in mouse spermatogenesis. Cell Rep. 2017;18(6):1383–94.

154. Pittman DL, Weinberg LR, Schimenti JC. Identification, characterization, and genetic mapping of Rad51d, a new mouse and human *RAD51*/*RecA*-related gene. Genomics. 1998;49(1):103–11.

155. Li XC, Bolcun-Filas E, Schimenti JC. Genetic evidence that synaptonemal complex axial elements govern recombination pathway choice in mice. Genetics. 2011;189(1):71–82.

156. Li S, Wang H, Jehi S, Li J, Liu S, Wang Z, et al. PIF1 helicase promotes break-induced replication in mammalian cells. EMBO J. 2021;40(8):e104509.

157. Lo HC, Hollingsworth NM. Using the semi-synthetic epitope system to identify direct substrates of the meiosis-specific budding yeast kinase, Mek1. Methods Mol Biol. 2011;745:135–49.

158. Goldstein AL, McCusker JH. Three new dominant drug resistance cassettes for gene disruption in *Saccharomyces cerevisiae*. Yeast. 1999;15(14):1541–53.

159. Longtine MS, McKenzie A, 3rd, Demarini DJ, Shah NG, Wach A, Brachat A, et al. Additional modules for versatile and economical PCR-based gene deletion and modification in *Saccharomyces cerevisiae*. Yeast. 1998;14(10):953–61.

160. Rothstein R. Targeting, disruption, replacement, and allele rescue: integrative DNA transformation in yeast. Methods Enzymol. 1991;194:281–301.

161. Boeke JD, Trueheart J, Natsoulis G, Fink GR. 5-Fluoroorotic acid as a selective agent in yeast molecular genetics. Methods Enzymol. 1987;154:164–75.

162. Falk JE, Chan AC, Hoffmann E, Hochwagen A. A Mec1- and PP4-dependent checkpoint couples centromere pairing to meiotic recombination. Dev Cell. 2010;19(4):599–611.

163. Grubb J, Brown MS, Bishop DK. Surface spreading and immunostaining of yeast chromosomes. J Vis Exp. 2015;(102):e53081.

164. Perkins DD. Biochemical mutants in the Smut fungus *Ustilago maydis*. Genetics. 1949;34(5):607–26.

165. Owens S, Tang S, Hunter N. Monitoring recombination during meiosis in budding yeast. Methods Enzymol. 2018;601:275–307.

166. Li H. Minimap2: pairwise alignment for nucleotide sequences. Bioinformatics. 2018;34(18):3094–100.

167. Lindenbaum P. JVarkit: java-based utilities for bioinformatics. 2016.

168. Sikorski. RS, Hieter P. A system of shuttle vectors and yeast host strains designed for efficient manipulation of the DNA in *Saccharomyces cerevisiae*. Genetics. 1989;122:19–27.

169. Parent, SA, Fenimore CM, Bostian, KA. Vector systems for the expression, analysis and cloning of DNA sequences in *S. cerevisiae*. Yeast. 1985;1:83–138.

